# Differential and Cooperative Immunomodulation by Saponin Adjuvants QS-21 and QS-7 in Human THP-1 Macrophages

**DOI:** 10.64898/2026.11.10.765708

**Authors:** Priyanka Talukdar, Emerson Smith, Jamie L. Inman, Harshini Mukundan

**Affiliations:** Lawrence Berkeley National Laboratory

## Abstract

QS-21, a triterpene glycoside saponin derived from *Quillaja saponaria*, is a potent immunostimulatory adjuvant incorporated in several FDA-approved vaccines. QS-7, a structurally related saponin fraction, has been reported to retain immunomodulatory activity with reduced hemolytic toxicity; however, its lower natural abundance has limited its systematic immunological characterization. To address this gap, we systematically compare responses to QS-21 and QS-7 in the THP-1 human macrophage model to resolve their distinct and cooperative contributions to antigen presenting cell (APC) activation. Following cytotoxicity profiling to establish safe and biologically relevant dosing, flow cytometry, proteomics, and transcriptomics were used to delineate the immunological programs engaged by each saponin alone and in combination. QS-7 alone induced minimal upregulation of cell surface activation markers and did not elicit a significant transcriptional signature, yet it drove dose-dependent secretion of a broad panel of pro-inflammatory cytokines and chemokines, revealing a dissociation between these two arms of macrophage activation. Strikingly, co-administration of QS-7 with QS-21 substantially potentiated macrophage activation across all parameters tested, amplifying surface marker upregulation, broadening cytokine and chemokine output to include IL-12p70 and IFN-γ, and markedly expanding the pro-inflammatory transcriptional and pathway landscape induced by QS-21, while concurrently suppressing counter-regulatory lipid-sensing pathways (PPAR, RAR, FXR/RXR) that constrain inflammatory output. Collectively, these findings establish QS-7 not merely as an attenuated analog of QS-21, but as a functionally distinct immunomodulator that recalibrates both the magnitude and regulatory tone of QS-21-driven macrophage activation. These findings show that structurally distinct saponins elicit distinct and complementary macrophage responses, providing a foundation for structure-function studies. Together with synthetic biology and data-driven design, this knowledge can guide the development of new-to-nature saponin adjuvants.

## Introduction

The development of effective vaccines against complex pathogens and emerging infectious diseases, has underscored the critical need for potent adjuvants capable of directing robust and durable adaptive immune responses. Among the most promising classes of natural immunostimulatory compounds are the triterpene glycoside saponins derived from the bark of the Chilean soapbark tree *Quillaja saponaria* Molina (family Quillajaceae) (1), collectively designated QS saponins based on their species of origin (*Q. saponaria*). These saponins have gained widespread recognition as vaccine adjuvants due to their ability to induce Th1-polarized immune responses and elicit cytotoxic T lymphocyte (CTL) activity against exogenous antigens (2–4)—properties that are particularly desirable for subunit and antigen-based vaccine platforms that are otherwise poorly immunogenic.

QS-21, the most extensively characterized fraction of the QS extract, is a mixture of two isomeric compounds each composed of four structural domains—a triterpene quillaic acid core, a branched trisaccharide unit, a linear tetrasaccharide chain, and a glycosylated acyl side chain (5). The lipophilic acyl side chain is critical for QS-21 immunostimulatory activity and may facilitate interactions with cellular membranes. QS-21 undergoes cholesterol-dependent endocytosis and subsequently accumulates in lysosomes, where it induces membrane destabilization and cathepsin B- and Syk-dependent activation of APCs (6,7). QS-21 serves as a key immunomodulatory component of the adjuvant system AS01, which is incorporated into two GlaxoSmithKline-developed human vaccines: Shingrix, approved for the prevention of herpes zoster (shingles), and Mosquirix, developed for malaria prophylaxis. Despite its clinical success and broad utility across vaccine platforms, QS-21 exhibits significant dose-dependent hemolysis and other toxicities that limits its use as a standalone adjuvant. Its incorporation into liposome-based formulations, such as AS01, improves tolerability (2,8). These limitations have driven interest in identifying structurally related saponin fractions that retain immunostimulatory potency while exhibiting improved tolerability profiles.

QS-7, another bioactive fraction isolated from the *Q. saponaria* extract, has similarly been reported to possess adjuvant activity, promoting strong CD8+ CTL responses and a balanced Th1/Th2 immune profile (9–11). Structurally, QS-7 differs markedly from QS-21: It lacks the hallmark long, chiral acyl side chain of QS-21 at the 4-O position of the fucopyranosyl residue that is typically linked to the C28 carboxyl group of quillaic acid—a feature replaced instead by a short acetyl group (Figure 1). Furthermore, QS-7 contains two additional monosaccharide residues within the C28 oligosaccharide domain relative to QS-21, conferring greater overall hydrophilicity. Given that the acyl side chain is central to QS-21’s membrane-intercalating and lysosome-destabilizing activity, this structural divergence raises the possibility that QS-7 engages APCs through a partially distinct activation mechanism, one potentially less reliant on membrane perturbation and more dependent on its extended, more hydrophilic oligosaccharide domain. Critically, QS-7 has been reported to exhibit reduced erythrocyte lytic activity and minimal systemic toxicity in murine models (2,11,12), suggesting a potentially more favorable safety profile. Although the adjuvanticity of QS-7 has been evaluated in the context of model protein antigens such as ovalbumin and viral antigens including HIV gp120 peptides (11), it has not progressed into any licensed vaccine formulation. This is largely attributable to challenges associated with its purification from the complex QS extract, its structural complexity, and the absence of large-scale clinical evaluation programs. Consequently, while QS-21 has been the subject of extensive immunological investigation over the past two decades, the cellular and molecular mechanisms underpinning QS-7’s immunomodulatory activity remain poorly defined.

**Figure 1:**
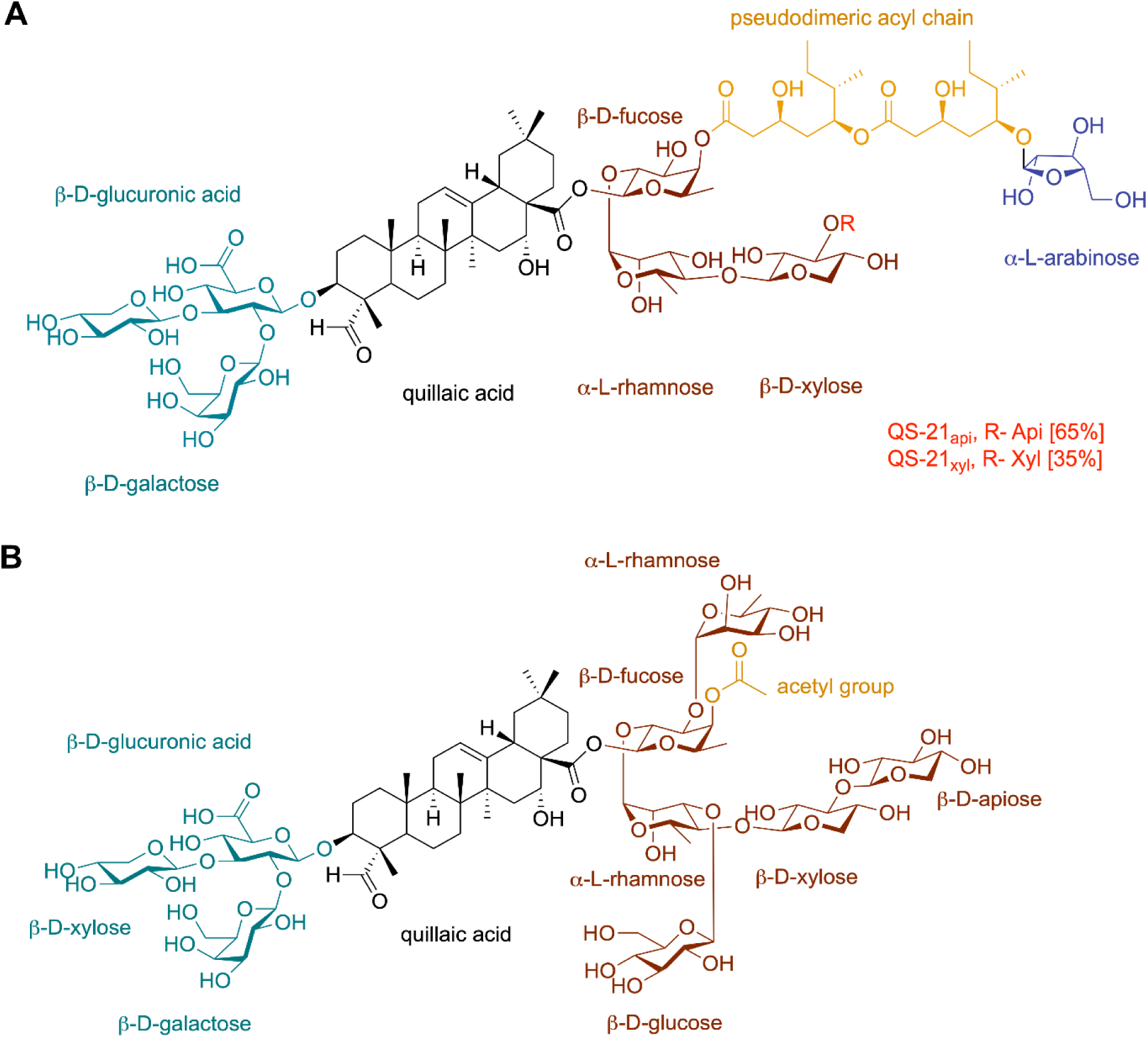
Chemical structures of saponin adjuvants QS-21 and QS-7. Schematic representations of the chemical structures of QS-21 (A) and QS-7 (B). Both molecules share a common triterpene quillaic acid core with a branched trisaccharide unit at the C3 position and a linear oligosaccharide chain at the C28 carboxyl group. Key structural differences are highlighted: QS-21 possesses a long, chiral acyl side chain at the 4-*O* position of the fucopyranosyl residue within the C28 oligosaccharide domain, whereas QS-7 bears a short acetyl group at this position. Additionally, QS-7 contains two extra monosaccharide residues within the C28 oligosaccharide domain relative to QS-21, contributing to its greater hydrophilicity. The indicated 65:35 QS-21-Api:QS-21-Xyl ratio is literature-reported (15,16) and was not determined for the preparation used in this study.

Macrophages, as professional APCs, are critical early mediators of adjuvant-driven innate immune activation. Their high adjuvant uptake capacity, coupled with their ability to activate dendritic cells (DCs) and orchestrate downstream antigen-specific B and T cell responses, positions them as key sentinels at the interface of innate and adaptive immunity (13,14). Adjuvant-mediated activation of macrophages typically culminates in upregulation of co-stimulatory surface molecules, secretion of pro-inflammatory cytokines and chemokines, and transcriptional reprogramming that collectively shape the magnitude and quality of the ensuing adaptive immune response. Characterizing these events at the cellular and molecular level is therefore essential not only for understanding adjuvant mechanisms of action, but also for the rational design of next-generation adjuvant systems with optimized efficacy and safety profiles.

Despite QS-21’s well-characterized immunological profile, established through more than two decades of research, a direct, side-by-side cellular and transcriptional comparison of QS-21 and QS-7 in human macrophages is entirely lacking. The mechanistic basis of QS-7’s immunomodulatory activity—at the level of APC activation, cytokine and chemokine induction, and transcriptional reprogramming—remains uncharacterized. This represents a critical gap, as such mechanistic insight is essential for evaluating QS-7’s potential as a viable adjuvant candidate, informing structure-activity relationships within the broader QS saponin family, and providing a scientific rationale for its advancement as a next-generation adjuvant with an improved tolerability margin relative to QS-21. Indeed, with the advancement in synthetic biology as a chassis for the production of saponins, and AI-enabled molecular re-design; a comprehensive characterization of structure-function relationships can drive rational new-to-nature adjuvant design in the future.

To address these questions, we employed the well-established THP-1 human macrophage model to perform a comprehensive, head-to-head immunological characterization of QS-21 and QS-7. Our findings reveal distinct but complementary immunological properties of QS-21 and QS-7. While QS-21 showed greater cytotoxicity and a stronger capacity to induce cell surface activation, QS-7 exhibited substantially lower cytotoxicity in epithelial cells while still promoting robust pro-inflammatory cytokine and chemokine secretion in macrophages. Notably, QS-7 enhanced both surface activation and inflammatory cytokine and chemokine responses when combined with QS-21, and transcriptomic profiling further demonstrated that their combination amplified inflammatory programs beyond those induced by QS-21 alone. Together, these findings indicate that the relationship between saponin structure, immunostimulatory activity, and cytotoxicity is more complex than can be captured by any single functional readout. Integrating complementary multi-omic approaches may therefore help identify structural features that govern these distinct properties and enable more rational design of saponin-based immunomodulators. More broadly, the complementary activity of individual saponin components suggests that rational combinations could provide a means to tune immunostimulatory activity while balancing potency and tolerability, with potential implications for the development of next-generation adjuvant formulations.

## Materials and Methods

### Cell Culture

THP-1 and A549 were purchased from ATCC (Manassas, VA). THP-1 were cultured in RPMI 1640 media (ATCC) supplemented with 10% FBS (Cytiva, Marlborough, MA), 100 U/mL penicillin-streptomycin (Cytiva) and 50 µM 2-mercaptoethanol (Gibco, Grand Island, NY). THP-1s were differentiated into macrophages with 100 ng/mL Phorbol 12-myristate 13-acetate (PMA) (Sigma Aldrich, St. Louis, MO) for 48 hours, followed by a 24 hour period of rest, before treatment with adjuvants. A549s were cultured in F-12K media (ATCC), supplemented with 10% FBS, 100 U/mL penicillin-streptomycin (Cytiva).

### Treatments

Cells were treated with QS-21 (GalaxoSmithKline; purity 96.4%, as determined by LC-MS) and QS-7 (Desert King international, Chula Vista, CA; Lot No. L181223PP05-2; purity 87.58%, as determined by LC-MS). Both saponins were dissolved in phosphate-buffered saline (PBS) at a stock concentration of 1 mg/mL, sterile filtered and stored in amber glass vials at -80C until use. Prior to treatment, stock solutions were diluted in cell culture medium to indicated final concentrations. Cells were treated with QS-21, QS-7, or their combination for the timepoints described in the experiment specific methods and figures prior to analysis. Cell culture medium without added saponins served as the vehicle control.

For the co-treatments, THP-1 macrophages were treated with QS-21 at a fixed final concentration of 5 μg/mL together with QS-7 at final concentrations of 2-20 μg/mL (2, 5, 10 and 20 μg/mL). QS-21 and QS-7 were each prepared from the 1 mg/mL stock solutions and diluted in cell culture medium. The two saponins were added to the cells simultaneously as separate additions rather than pre-mixed. Concentrations refer to the final concentration of each saponin in the cell culture well. Co-treatment groups were compared with the QS-21 (5 μg/mL) alone condition.

### PrestoBlue Assay

PrestoBlue HS assay (Thermo Fisher Scientific, Waltham, MA) was performed to compare the cytotoxic effects of QS-21 and QS-7 in A549 and THP-1 cells, following the manufacturer’s protocol. Briefly, THP-1–derived macrophages and A549 cells were treated with the adjuvants at concentrations ranging from 0.1 μg/mL to 20 μg/mL for 24 hours. Cells were then incubated with one-tenth the volume of PrestoBlue HS, and absorbance was measured after 30 minutes.

### Flow Cytometry

Following treatment with adjuvants for 24 hours, supernatants were collected and stored at −80 °C for luminex analysis and adherent macrophages were washed with 1x PBS (ThermoFisher Scientific). Cells were next treated with 1x Versene (Gibco) for 25 minutes at 37 ℃, as it provides non-enzymatic cell detachment, helping avoid enzymatic cleavage of surface epitopes. All treatment groups were detached and processed in parallel using identical handling, times and volumes. Remaining adherent cells were gently scraped from the plate, washed with 1x PBS and stained for viability (Live/Dead Fixable Red, Thermo Fisher Scientific) and cell surface activation markers according to standardized protocol. The antibody panel included CD86 (Clone 2331 (FUN-1) (RUO), Waters Biosciences, San Jose, CA), CD80 (Clone 2D10.4, ThermoFisher Scientific), CD40 (Clone 5C3, ThermoFisher Scientific) and HLA-DR (Clone eBM2a, ThermoFisher Scientific). Flow cytometry data was acquired with Attune Cytpix. Compensation controls were prepared using UltraComp eBeads Compensation Beads (ThermoFisher Scientific) for antibody fluorophores and the ArC Amine Reactive Compensation Bead Kit (ThermoFisher Scientific) for the live/dead stain, following the manufacturers’ protocols. Data analyzed with FlowJo v10 (Waters Biosciences). Cells were first gated based on forward scatter (FSC) and side scatter (SSC), followed by gating of viable cells using the Live/Dead Fixable Red viability dye (Supplementary Figure 1). Surface marker expression (CD80, CD86, CD40, and HLA-DR) was then gated using the corresponding isotype controls. Marker expression was quantified as geometric mean fluorescence intensity (gMFI).

### Luminex

THP-1 derived macrophages were treated with QS-21 and QS-7 for 24 hours and cytokines and chemokines produced by the macrophages were measured by Luminex using Human Luminex Discovery Assay (Catalog-LXSAHM, R&D Systems, Minneapolis, MN) according to the manufacturer’s protocol. The analytes studied and their limit of detection (LOD) values are TNFα (1.2 pg/mL), IL-6 (1.7 pg/mL), IL-1β (0.8 pg/mL), IL-18 (1.93 pg/mL), GM-CSF (4.1 pg/mL), CCL2 (9.9 pg/mL), CXCL1 (5.3 pg/mL), CXCL10 (1.18 pg/mL), IL-12p70 (20.2 pg/mL), IFNγ (0.22 pg/mL), IL-27 (2.7 pg/mL) and IL-10 (1.6 pg/mL). A 1:2 dilution was used for the samples. Data was acquired using the MAGPIX instrument and analyzed with xPONENT software. Cytokine concentrations in culture supernatants were quantified and reported in pg/mL. Analytes that did not give detectable values are represented as half the value of the LOD solely for visualization purposes with the LOD represented as a dashed line. Analytes that were above the LOD but below the lower limit of quantification (LLOQ) have been highlighted in the manuscript results section. To account for differences in viable-cell numbers between treatment conditions, cytokine concentrations were normalized to the total number of viable cells in the corresponding well. Following supernatant collection, cells from each well were assessed by flow cytometry using live/dead staining. The total viable-cell count per well was calculated from the flow-cytometric assessment, accounting for the total volume of the cell suspension prepared for analysis. For each well, the measured cytokine concentration (pg/mL) was divided by the estimated total number of viable cells and multiplied by (10=6). Normalized values are reported as Luminex-measured cytokine concentration per (10^6) viable cells.

### RNA Sequencing and Pathway Analysis

THP-1 cells post differentiation into macrophages were treated with 5 ug/mL of QS-21 or 5 ug/mL QS-7 or 5 ug/mL each of QS-21 and QS-7 combined for 5 hours. RNA was isolated using Qiagen RNEASY mini kit (Redwood City, CA) according to the manufacturer’s instructions and samples were submitted to University of California Los Angeles (UCLA) Technology Center for Genomics and Bioinformatics for sequencing and Next-generation sequencing (NGS) partial data analysis. Total RNA quality was checked with NanoDrop and Tapestation and sequencing done with Novaseq X Plus (Supplementary Table 1). Raw gene count data generated from RNA sequencing were analyzed in R version 4.5.2 (R Foundation for Statistical Computing, Vienna, Austria) using the edgeR and limma Bioconductor packages (17). Samples comprised four experimental conditions, including one control condition and three treatment conditions, each represented by three matched biological replicates collected on independent experimental days.

Raw gene counts were imported into an edgeR DGEList object, and normalization factors were calculated using the trimmed mean of M-values (TMM) method to account for differences in library composition. Lowly expressed genes were removed using the filterByExpr() function with default parameters, after which normalization factors were recalculated. Normalized count data were transformed to log2 counts per million (logCPM) using the voom function in the limma package, which estimates the mean–variance relationship of the log-count data and assigns precision weights to each observation for linear modeling.

Differential gene expression analysis was performed using the limma-voom pipeline. Because all four experimental conditions (control, QS-21, QS-7, and QS-21+QS-7 combination) were evaluated across the same three independent experimental days, the statistical model accounted for both treatment condition and experimental day to account for the matched experimental design and reduce the influence of day-to-day variability on treatment comparisons. Raw gene counts were imported into edgeR, normalized using the trimmed mean of M-values (TMM) method, and filtered to remove genes with insufficient expression. The filtered counts were then transformed using voom, which estimates the mean–variance relationship of log-counts and generates precision weights for use in linear modeling. Gene-wise linear models were fitted to estimate treatment-associated expression differences, and empirical Bayes moderation was applied to improve the stability of variance estimates, particularly given the limited number of biological replicates. Pairwise contrasts compared each treatment condition with the control. Differential expression statistics were extracted using topTable(), with P-values adjusted for multiple testing using the Benjamini–Hochberg method to control the false discovery rate (FDR). Genes were considered significantly differentially expressed if they exhibited an adjusted P-value (FDR) < 0.05 and an absolute log2 fold change (|logFC|) ≥ 2. Gene annotations, including Ensembl gene identifiers and gene symbols, were obtained from the annotation accompanying the original raw count file to maintain consistency with the reference genome and annotation release used during read quantification.

Volcano plots were generated using the EnhancedVolcano R package (18). Euler plots illustrating the overlap of differentially expressed genes (DEGs) among the comparisons were generated using the R package eulerr (19). Gene Ontology (GO) enrichment analysis was performed using the topGO R package. Genes with an adjusted P-value (Benjamini–Hochberg) below 0.05 were ranked according to the absolute value of their log fold change (logFC), and up to 1,000 genes with the largest absolute fold changes were selected for analysis. All genes tested in the differential expression analysis and successfully mapped to Entrez Gene IDs were used as the background gene universe. Ensembl gene identifiers were converted to Entrez Gene IDs using the Bioconductor packages AnnotationDbi and org.Hs.eg.db. GO annotations were retrieved, and analysis was restricted to the Biological Process (BP) ontology. Enrichment was assessed using Fisher’s exact test with the weight01 algorithm and a minimum GO term size of 10 genes. P-values were adjusted for multiple testing using the Benjamini–Hochberg method, with an FDR below 0.05 considered statistically significant. Fold enrichment was calculated as the ratio of observed to expected selected genes for each GO term.

Differentially expressed genes obtained from each comparison of interest were analyzed using Ingenuity Pathway Analysis (IPA; QIAGEN Inc.) to identify enriched canonical pathways. Ensembl gene identifiers were submitted to IPA and mapped using the IPA knowledge repository. Pathway analysis was performed using the mapped genes, with pathway significance evaluated based on Benjamini–Hochberg-adjusted P-values (FDR) and activation z-scores. Pathways with an adjusted P-value (FDR) < 0.05 and an absolute z-score ≥ 2 were considered significant. The results were subsequently used in R to plot significantly upregulated pathways using the ggplot2 package (20).

### Statistics

Statistical analyses were performed using GraphPad Prism version 11.0.0 (GraphPad Software, San Diego, CA). Data distributions were assessed for normality using the Shapiro–Wilk test. For flow cytometry and Luminex endpoints, repeated-measures one-way ANOVA followed by Dunnett’s multiple-comparisons test was used for normally distributed data, and non-normally distributed data were analyzed using the Friedman test followed by Dunn’s multiple-comparisons test, as specified in the corresponding figure legends. The Geisser–Greenhouse correction was applied to account for violations of the sphericity assumption, where appropriate. Data are presented as mean ± standard error of the mean (SEM), with the number of independent biological replicates indicated in each figure legend. Statistical significance was defined as P < 0.05. For RNA-sequencing analyses, differential expression P-values were adjusted for multiple testing using the Benjamini–Hochberg method, with an FDR < 0.05 and an absolute log2 fold change ≥ 2 used to define significantly differentially expressed genes.

## Results

### Differential Cytotoxicity Profiles of QS-21 and QS-7 in THP-1 Macrophages and A549 Epithelial Cells

Although QS-7 is reported to be less hemolytically toxic than QS-21, a systematic, dose-dependent cytotoxicity profile for either compound in human cell lines has not been established. To address this, we first sought to define the cytotoxic dose range of QS-21 and QS-7 in two cell line models —THP-1 monocyte-derived macrophages and A549 lung epithelial cells—using the PrestoBlue HS cell viability assay, which measures cellular metabolic activity as a proxy for viability. The two models were selected to capture potential differences in cytotoxic sensitivity between immune cells and lung epithelial cells, thereby providing a broader assessment of cellular tolerance to the saponins. Both compounds were evaluated across a dose range of 0.1–20 μg/mL to comprehensively characterize their cytotoxic thresholds across both cell types.

A549 epithelial cells tolerated increasing concentrations of QS-21 well up to 10 μg/mL, with a notable decrease in metabolic activity observed only at 20 μg/mL (Figure 2A). THP-1 macrophages, however, exhibited greater sensitivity, with QS-21 inducing a pronounced dose-dependent reduction in cellular metabolic activity consistent with previously reported toxicity data, falling below 10% at the highest tested concentration of 20 μg/mL (Figure 2A). In contrast, QS-7 demonstrated markedly reduced cytotoxicity in A549 cells, with greater than 90% of cells remaining metabolically active even at the maximum tested dose of 20 μg/mL (Figure 2B), confirming its superior tolerability in epithelial cells. QS-7 similarly induced a dose-dependent decline in THP-1 macrophage metabolic activity; however, at the highest tested concentration of 20 μg/mL, metabolic activity was higher than that observed with QS-21 at the same concentration (Figure 2B). Taken together, these data establish the differential cytotoxicity profiles of QS-21 and QS-7 across two distinct cell types and informed the selection of working concentrations for subsequent immunological characterization experiments.

**Figure 2:**
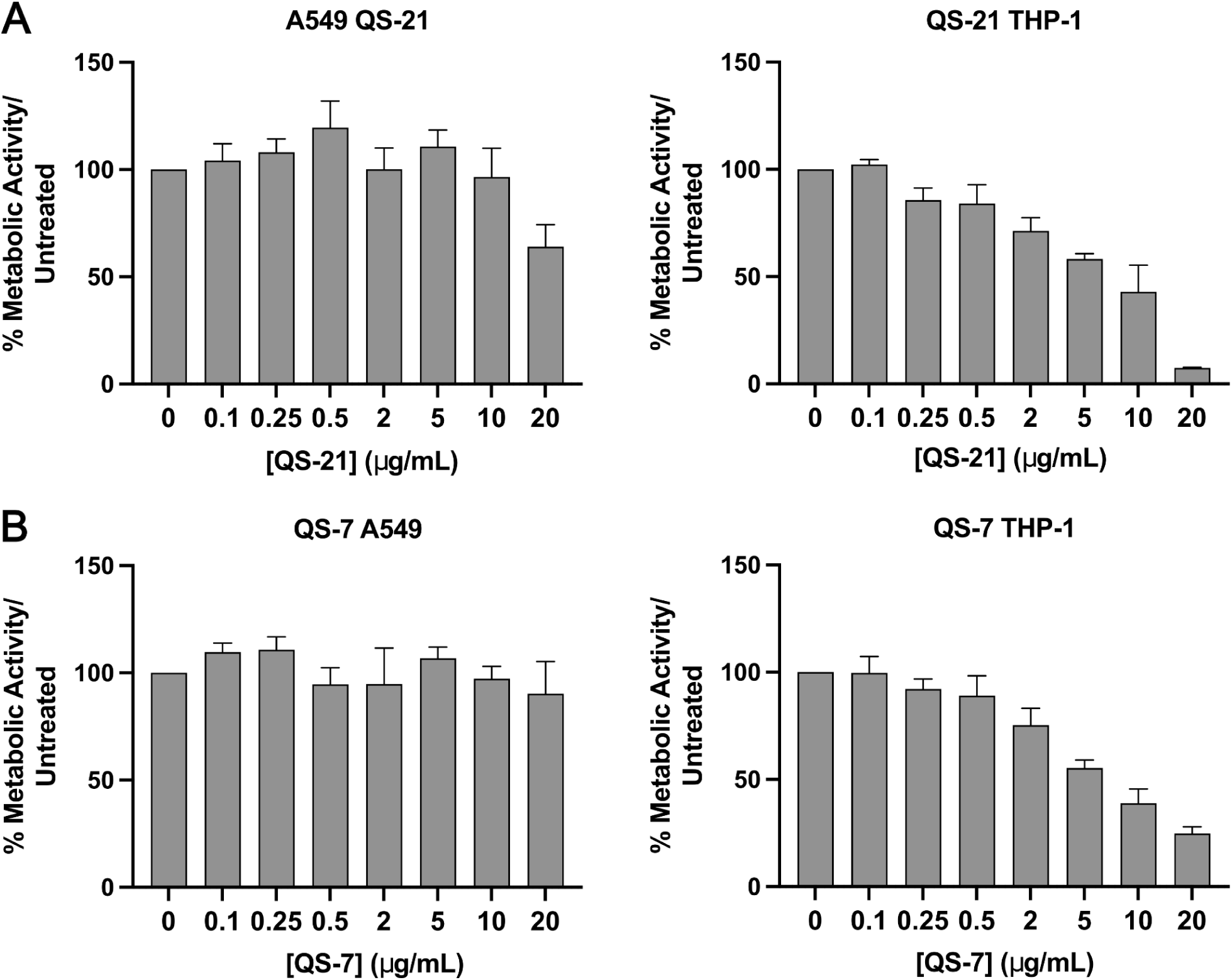
Cytotoxicity profiles for QS-21 and QS-7 in A549 epithelial cells and THP-1 macrophages. Cellular metabolic activity, measured as a proxy for viability using the PrestoBlue HS assay, in A549 lung epithelial cells and THP-1 monocyte-derived macrophages treated with increasing concentrations of QS-21 (A) or QS-7 (B) for 24 hours. (A) A549 cells tolerate QS-21 well up to 10 μg/mL, with a notable decline in metabolic activity observed at 20 μg/mL. THP-1 macrophages exhibit greater sensitivity, showing a pronounced dose-dependent reduction in metabolic activity across the tested concentration range. (B) QS-7 does not induce significant cytotoxicity in A549 cells across the full dose range tested (0.1–20 μg/mL), whereas THP-1 macrophages again display a dose-dependent decline in metabolic activity with increasing QS-7 concentrations. Data are presented as mean ± SEM of n=3 independent biological replicates.

### QS-7 Potentiates QS-21-Mediated Macrophage Surface Activation Despite Limited Independent Activity

QS-21 has been reported to upregulate the expression of HLA-DR and CD86 in human monocytes, myeloid DCs and monocyte-derived DCs (moDCs) (6,21), however, the capacity of QS-7 to modulate cell surface activation marker expression on macrophages remains entirely uncharacterized. To address this, we evaluated the effects of QS-21 (2–10 μg/mL) and QS-7 (2–20 μg/mL) on the expression of the co-stimulatory molecules CD80, CD86, and CD40, and the antigen-presentation molecule HLA-DR in THP-1 macrophages by flow cytometry. QS-21 induced significant upregulation of CD80 and HLA-DR at 10 μg/mL, and CD86 expression was significantly elevated from 5 μg/mL onwards (Figure 3A, Supplementary Table 2). In contrast, QS-7 across its entire tested dose range failed to induce any significant change in the expression of any of the four activation markers examined (Figure 3B, Supplementary Table 2), indicating that QS-7 alone is insufficient to drive THP-1 macrophage surface activation at the concentrations tested.

**Figure 3:**
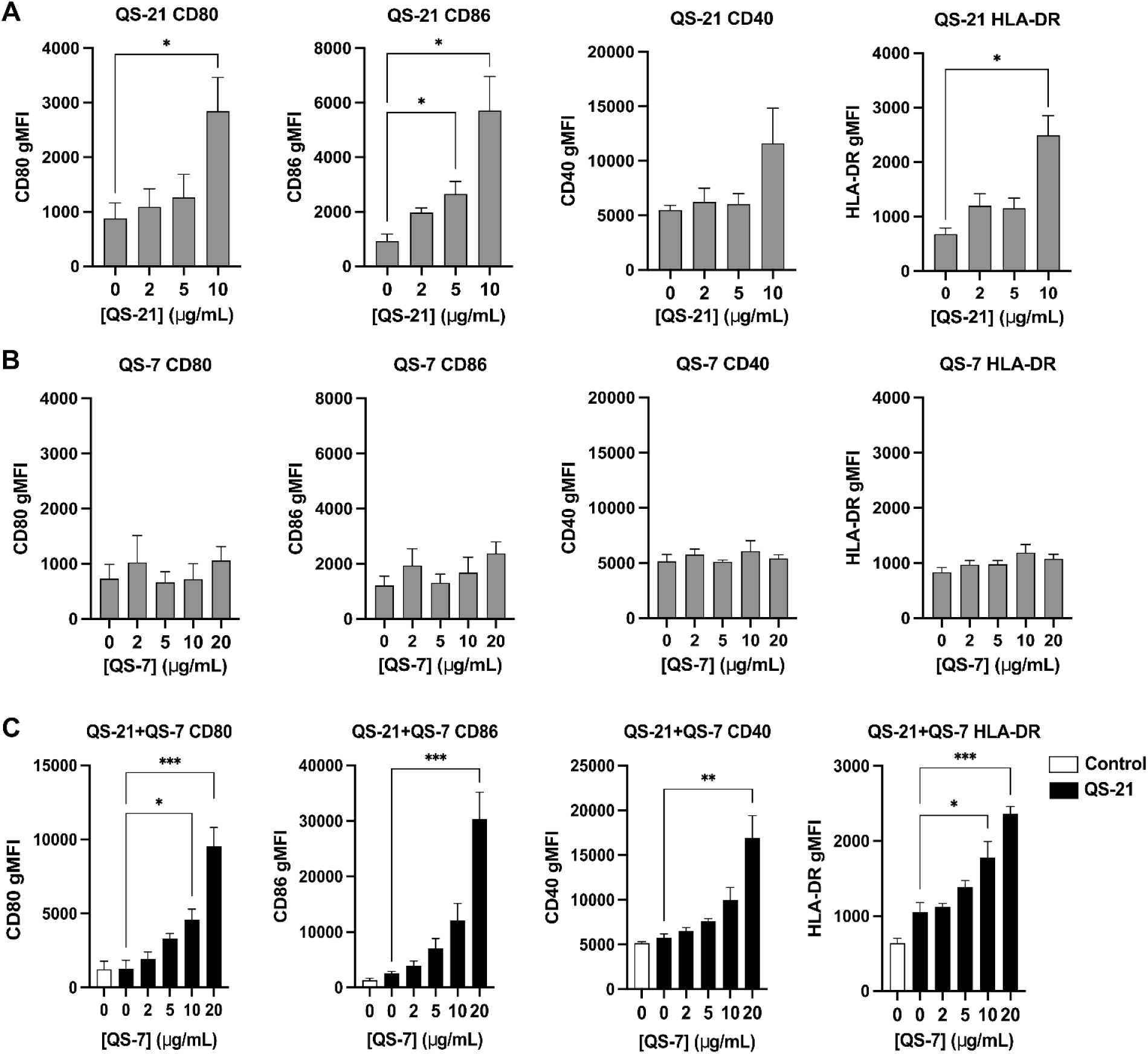
Differential modulation of cell surface activation marker expression by QS-21 and QS-7 in THP-1 macrophages. Flow cytometric quantification of co-stimulatory and antigen-presentation molecules CD80, CD86, CD40, and HLA-DR on THP-1 macrophages following treatment with increasing concentrations of QS-21 or QS-7 alone, or QS-7 in combination with a fixed concentration of QS-21 (5 μg/mL). (A) QS-21 induces significant, dose-dependent upregulation of CD80, CD86, and HLA-DR in THP-1 macrophages. Data are presented as mean ± SEM of n = 5 independent biological replicates. (B) Treatment with increasing concentrations of QS-7 alone does not result in significant upregulation of any of the cell surface activation markers examined. Data are presented as mean ± SEM of n = 5 independent biological replicates. (C) Although QS-7 alone is insufficient to drive cell surface activation marker expression, co-treatment with increasing concentrations of QS-7 and a fixed sub-maximal concentration of QS-21 (5 μg/mL) results in a dose-dependent enhancement of CD80, CD86, CD40 and HLA-DR expression beyond levels elicited by QS-21 alone. Data are presented as mean ± SEM of n = 6 independent biological replicates. Statistical significance was assessed using repeated measures one-way ANOVA with Dunnett’s multiple comparisons test for normally distributed data and Dunn’s multiple comparisons test otherwise. For panels A and B, comparisons were made against the untreated control condition; for panel C, comparisons were made against the QS-21 alone condition. *p < 0.05, **p < 0.01, ***p < 0.001.

Given that QS-7 is not employed as a standalone adjuvant but may be present as a constituent of crude Quil-A fractions used in ISCOM formulations, and that prior murine studies have reported synergistic immune augmentation upon co-administration of QS-7 with QS-21 (12), we then investigated whether QS-7 could potentiate the immunostimulatory activity of QS-21 in THP-1 macrophages. Cells were treated with a fixed sub-maximal concentration of QS-21 (5 μg/mL) in combination with increasing concentrations of QS-7 (2–20 μg/mL). All comparisons were made against treatment with QS-21 (5 μg/mL), allowing any enhancement in activation marker expression to be specifically attributed to the contribution of QS-7. Co-treatment resulted in a dose-dependent enhancement of activation marker expression relative to QS-21 alone, with significant upregulation of CD80 and HLA-DR observed from 10 μg/mL QS-7 onwards, and significant increases in CD86 and CD40 expression were detected only at the highest tested QS-7 concentration (20 μg/mL) (Figure 3C, Supplementary Table 2). These findings demonstrate that, while QS-7 lacked an independent capacity to induce macrophage surface activation under our experimental conditions, it enhanced QS-21-mediated upregulation of activation and antigen-presentation markers, consistent with increased macrophage activation and antigen-presenting capacity. This suggests a cooperative immunomodulatory relationship between the two saponin fractions, with potential implications for adjuvant combination strategies.

### Both QS-21 and QS-7 Promote Pro-inflammatory Cytokine and Chemokine Production in THP-1 Macrophages with Distinct Dose-Dependent Potencies

QS-21 has been previously reported to induce the production of TNF, IL-6 and IL-8 in human moDCs (6), and to promote IL-6 secretion and inflammasome activation when combined with MPLA in murine DCs and macrophages, resulting in downstream IL-1β production (22). However, whether QS-7 shares this capacity to drive pro-inflammatory cytokine and chemokine production in human macrophages remains unknown. To enable a direct, side-by-side comparison and establish a functional reference benchmark for QS-7, both adjuvants were evaluated in parallel, given that a comprehensive cytokine and chemokine induction profile of QS-21 in THP-1 macrophages has not been previously defined. THP-1 macrophages were therefore stimulated with increasing concentrations of QS-21 (2–10 μg/mL) and QS-7 (2–20 μg/mL), and the secretion of a panel of mediators—TNFα, IL-6, IL-1β, IL-12p70, IFNγ, IL-18, IL-10, IL-27, GM-CSF, CXCL1, CXCL10 and CCL2—was quantified.

QS-21 induced robust, dose-dependent secretion of several inflammatory cytokines and chemokines in THP-1 macrophages. Significant upregulation of IL-6 and IL-1β was observed from the lowest tested concentration of 2 μg/mL, while TNFα, IL-18, CXCL1, CXCL10, and CCL2 reached significance from 5 μg/mL onwards. GM-CSF secretion was significant only at the highest concentration of 10 μg/mL QS-21 (Figure 4A). Notably, and in contrast to its inability to independently upregulate cell surface activation markers, QS-7 demonstrated a comparable capacity to drive pro-inflammatory cytokine and chemokine secretion in THP-1 macrophages. QS-7 similarly elicited dose-dependent responses, with significant upregulation of CXCL1 and CCL2 from 5 μg/mL, IL-6 and IL-1β from 10 μg/mL, and TNFα, IL-18, and GM-CSF reaching significance at the highest tested concentration of 20 μg/mL (Figure 4B). CXCL10 levels showed increasing trends of secretion with increasing concentrations of QS-7, although no statistically significant differences were observed. In both the cases where cells were treated with QS-21 or QS-7 independently. In both the cases where cells were treated with QS-21 or QS-7 independently, IL-12p70, IFNγ, IL-10, and IL-27 were not consistently detected across all biological replicates with levels across biological replicates remaining largely variable and below reliable detection thresholds (Supplementary Figure 2). Taken together, these results indicate that QS-7-mediated macrophage activation is not uniformly reflected across all measured parameters of immune activation, and that cytokine and chemokine secretion and cell surface activation marker upregulation might represent distinct, independently regulated facets of the macrophage response to QS-7. Furthermore, while QS-7 demonstrated attenuated potency relative to QS-21 at equivalent doses, the qualitative overlap in the spectrum of pro-inflammatory mediators induced by both adjuvants suggests engagement of shared, albeit differentially activated, downstream inflammatory signaling pathways.

**Figure 4:**
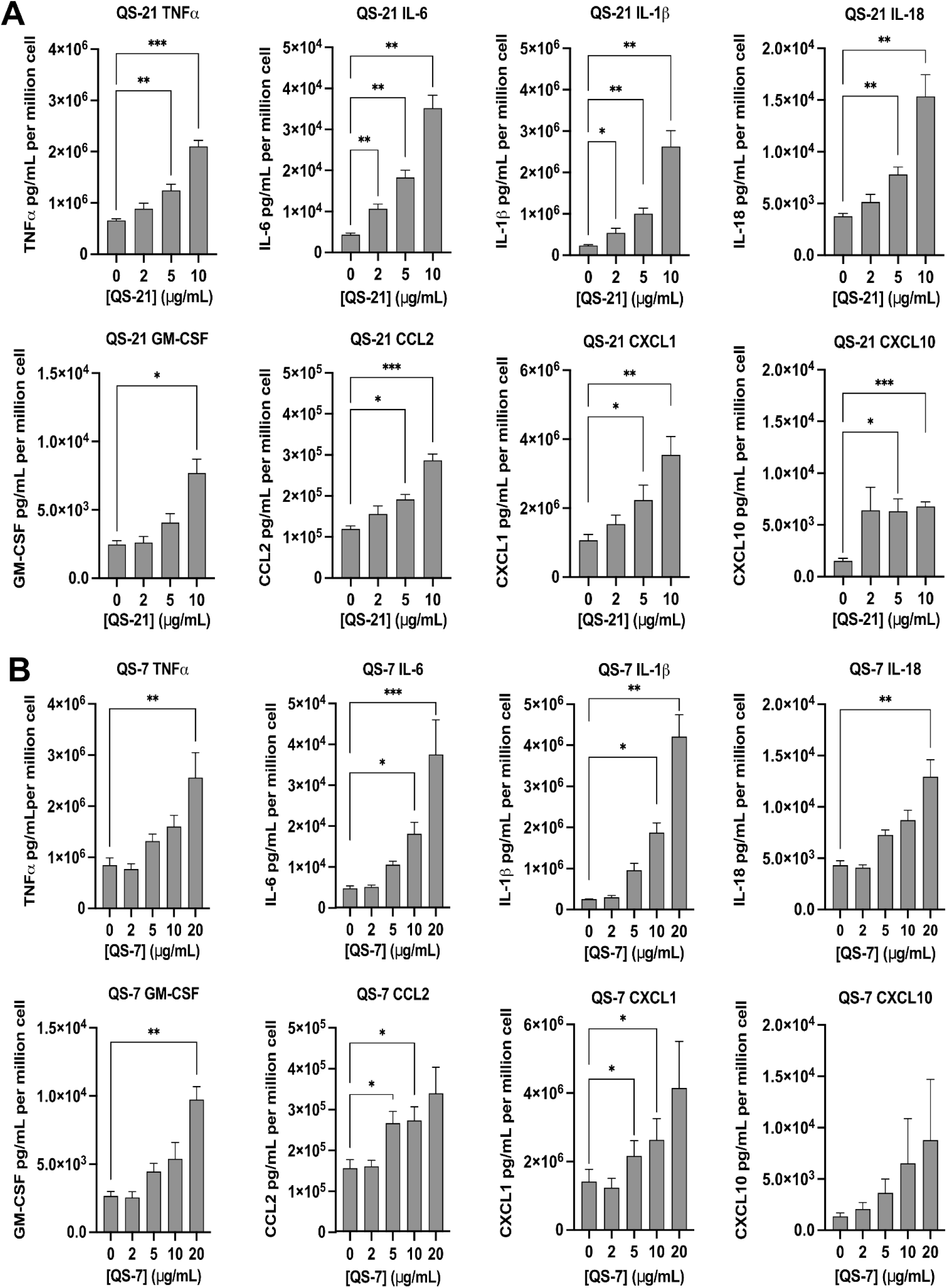
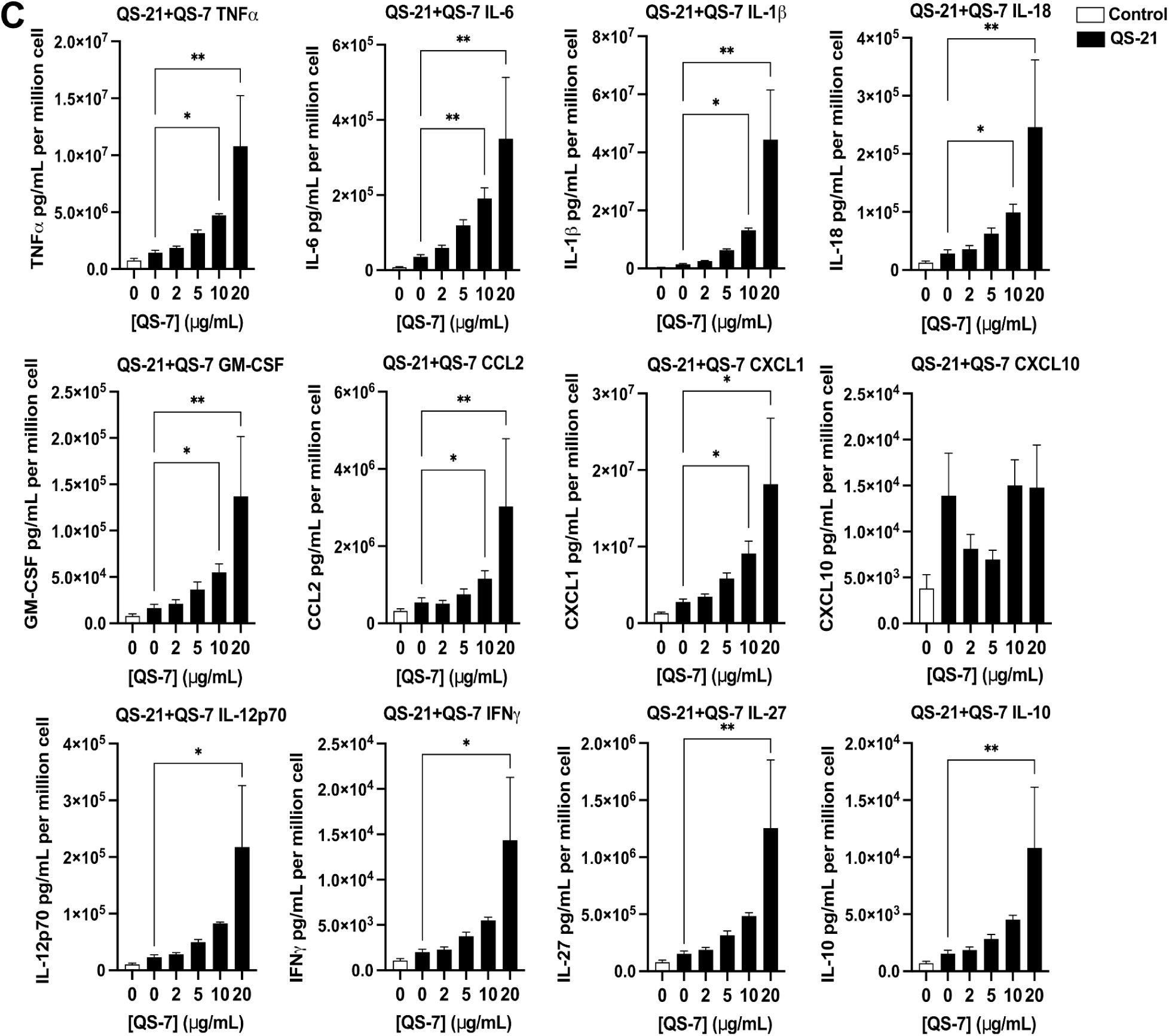
Both QS-21 and QS-7 promote pro-inflammatory cytokine and chemokine secretion in THP-1 macrophages, with QS-7 potentiating the response elicited by QS-21. Luminex multiplex quantification of cytokine and chemokine secretion in THP-1 macrophages treated with increasing concentrations of QS-21 or QS-7 alone, or QS-7 in combination with a fixed sub-maximal concentration of QS-21 (5 μg/mL). (A) QS-21 induces significant, dose-dependent upregulation of pro-inflammatory cytokines TNFα, IL-6, IL-1β, and IL-18, the hematopoietic growth factor GM-CSF, and chemokines CCL2, CXCL1, and CXCL10 in THP-1 macrophages. Data are presented as mean ± SEM of n = 5 independent biological replicates. (B) QS-7 similarly induces dose-dependent upregulation of TNFα, IL-6, IL-1β, IL-18, GM-CSF, CCL2, and CXCL1, albeit at comparatively higher concentrations relative to QS-21 to achieve significance. Data are presented as mean ± SEM of n = 5 independent biological replicates. (C) Mirroring the pattern observed for cell surface activation markers, co-treatment with increasing concentrations of QS-7 alongside a fixed sub-maximal concentration of QS-21 (5 μg/mL) results in a dose-dependent enhancement of cytokine and chemokine secretion beyond levels elicited by QS-21 alone. Data are presented as mean ± SEM of n = 6 independent biological replicates. Statistical significance was assessed using repeated measures one-way ANOVA with Dunnett’s multiple comparisons test for normally distributed data and Dunn’s multiple comparisons test otherwise. For panels A and B, comparisons were made against the untreated control condition; for panel C, comparisons were made against the QS-21 alone condition. *p < 0.05, **p < 0.01, ***p < 0.001.

Given the potentiating effect of QS-7 on QS-21-mediated surface activation marker expression, we then investigated whether the adjuvant could similarly amplify the cytokine and chemokine secretory response elicited by QS-21. Mirroring the co-treatment strategy used for surface marker analysis, THP-1 macrophages were treated with a fixed sub-maximal concentration of QS-21 (5 μg/mL) and increasing concentrations of QS-7 (2–20 μg/mL). QS-21-alone served as the reference condition to determine whether QS-7 provided additional immunostimulatory activity. Co-treatment resulted in a robust, dose-dependent enhancement of cytokine and chemokine secretion, with significant upregulation of TNFα, IL-6, GM-CSF, CCL2, IL-1β, CXCL1, and IL-18 observed from 10 μg/mL QS-7 onwards (Figure 4C). For CXCL10, highest levels were observed with 10 and 20 µg/mL QS-7 co-treatment, although no consistent enhancement over QS-21 alone was evident. In addition, significant increases in IL-12p70, IFNγ, IL-27, and IL-10 levels were observed with co-treatment of 20 ug/mL QS-7 in comparison to QS-21 alone (Figure 4C). These cytokines were not consistently induced by QS-21 or QS-7 alone, indicating a cooperative effect of the combined treatment. IL-12p70 and IFNγ are hallmark drivers of Th1 polarization and CTL responses (23–25)—immune programs that are central to the adjuvant activity attributed to QS saponins. However, the levels of these 2 cytokines were above the LOD but below the LLOQ, indicating detectable signals that could not be reliably quantified. The concurrent reproducible induction of IL-27 and IL-10 suggests the involvement of cytokines associated with immunomodulatory processes (26,27), potentially reflecting the engagement of pathways associated with the regulation of inflammatory responses. Collectively, these findings demonstrate that QS-7 not only amplifies the pro-inflammatory secretory response of QS-21 but qualitatively expands the cytokine milieu upon co-treatment, engaging immunological programs that are not reliably elicited by either adjuvant independently.

### QS-7 Amplifies the Transcriptional and Pathway Activation Program Induced by QS-21 in THP-1 Macrophages

The effects of saponin adjuvants on the gene expression profiles of immune cells remain largely uncharacterized. In one of the few existing studies, microarray analysis on RNA from QS-21 stimulated human moDCS demonstrated upregulation of genes associated with inflammatory cytokine and chemokine responses, as well as transcription factor networks governing genes induced by cellular stress, cytokines, and infectious agents (6). However, a comprehensive transcriptional analysis of saponin adjuvant activity in human macrophages, and particularly any comparison between QS-21 and QS-7, is entirely absent from the literature. To identify the immune response genes and pathways regulated by each adjuvant individually, and to characterize the transcriptional programs engaged upon their co-administration, RNA sequencing was performed on THP-1 macrophages treated with 5 μg/mL QS-21, 5 μg/mL QS-7, or the combination of QS-21 and QS-7 (QS-21+QS-7, both at 5 μg/mL) for 5 hours (Supplementary Figure 3). Genes with an adjusted P-value (FDR) < 0.05 and an absolute log2 fold change (|log2FC|) ≥ 2 were considered significantly differentially expressed (Supplementary Tables 3-5).

QS-21 treatment resulted in significant upregulation of 16 genes and downregulation of 2 genes, while QS-7 treatment alone did not yield a significant number of differentially expressed genes meeting the FDR threshold of < 0.05 (Figure 5A), consistent with its attenuated effect on surface activation marker expression observed above. Strikingly, the greatest transcriptional response was observed in macrophages treated with the QS-21+QS-7 combination, which resulted in significant upregulation of 61 genes and downregulation of 8 genes (Figure 5A), demonstrating that QS-7 substantially amplifies the transcriptional program initiated by QS-21. Given the absence of significant independent transcriptional activity by QS-7, subsequent analyses focused on delineating the differences between the transcriptional signatures of QS-21 and QS-21+QS-7. Of the 16 genes significantly upregulated by QS-21, 14 were shared with the QS-21+QS-7 condition, and 1 of the 2 genes downregulated by QS-21 was similarly downregulated in the combination condition (Figure 5B). Genes associated with inflammatory response (*TNF*, *IL-6*, *IL-23A*, *IL-36G*), chemotaxis and leukocyte recruitment (*CSF3*, *CCL3*, *CCL4*, *CXCL2*, *CXCL10*), transcriptional regulation (*IRF4*, *NFKB1A*) and APC maturation and migration (*CD80*, *CD83*, *CCR7*) were expressed at comparatively higher levels in QS-21+QS-7-treated macrophages relative to QS-21 alone (Figure 5C). A quick comparison of the differentially expressed genes in QS-21+QS-7-treated macrophages in comparison to QS-21 alone, identified *IL-6*, *IL-23A* and *HCAR2* to be significantly upregulated (Supplementary Table 6). Gene Ontology (GO) enrichment analysis identified 33 biological processes significantly enriched in the QS-21+QS-7 condition (Supplementary Tables 7-8), These processes included pathways associated with immune responses, T-cell costimulation, and positive regulation of type II interferon (IFNγ) production. The top 12 enriched pathways were predominantly associated with immune cell responses, cell migration, and inflammatory responses (Figure 5D), collectively indicating a broader and more robust pro-inflammatory and immunostimulatory transcriptional program compared with QS-21 alone.

**Figure 5:**
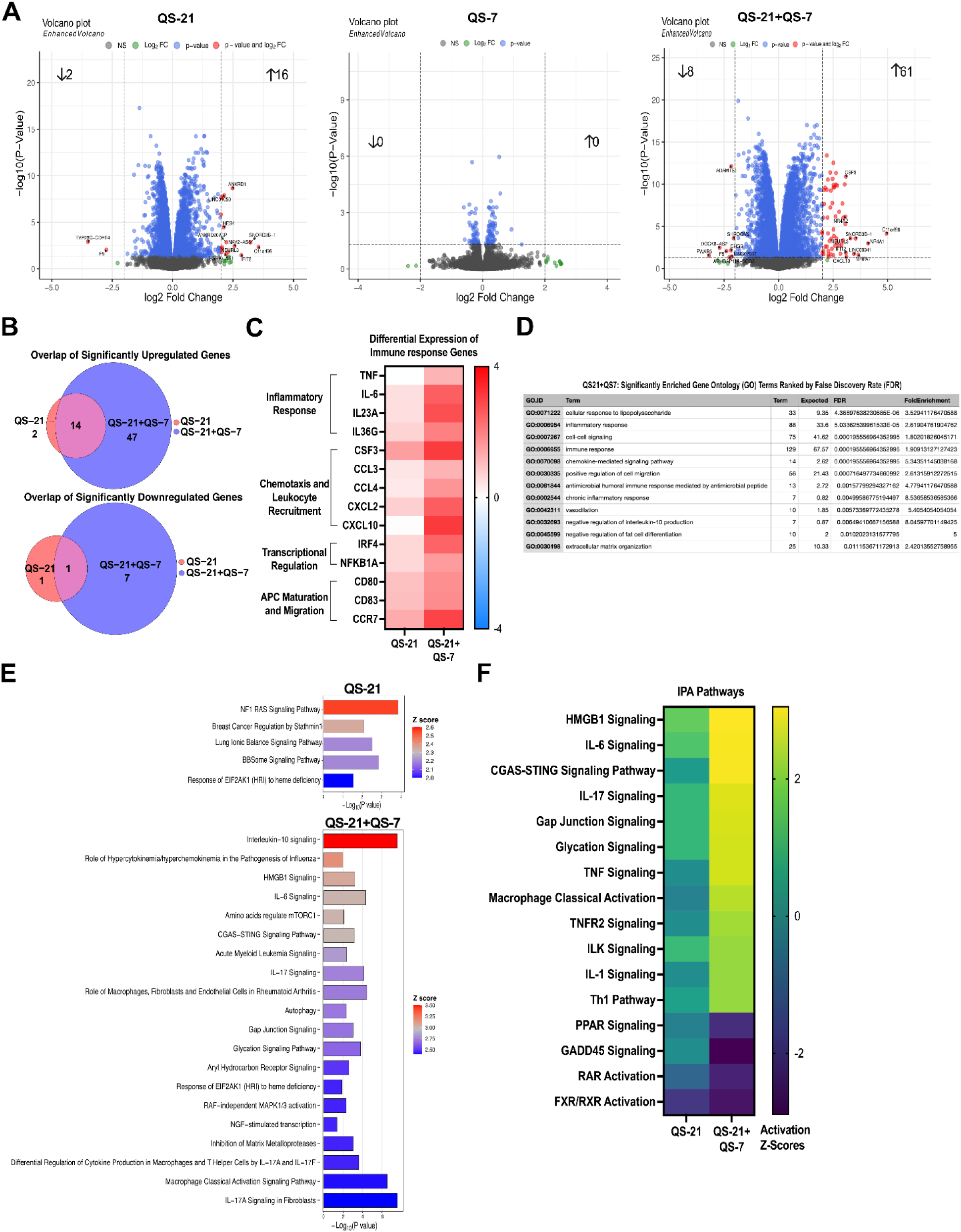
Transcriptional profiling reveals QS-7-mediated amplification of QS-21-induced pro-inflammatory gene networks in THP-1 macrophages. RNA sequencing was performed on THP-1 macrophages treated with QS-21 (5 μg/mL), QS-7 (5 μg/mL), or the combination of QS-21 (5 μg/mL) and QS-7 (5 μg/mL) (QS-21+QS-7). Genes with an adjusted p-value (FDR) < 0.05 and absolute log2 fold change (|log2FC|) ≥ 2 were considered significantly differentially expressed. (A) Volcano plots depicting the transcriptome-wide differential gene expression in THP-1 macrophages treated with QS-21, QS-7, or QS-21+QS-7 relative to untreated controls. The number of significantly upregulated and downregulated genes are highlighted. (B) Euler diagrams illustrating the overlap in significantly upregulated and downregulated genes between QS-21 and QS-21+QS-7 conditions, demonstrating that the combination treatment largely expands upon rather than redirects the transcriptional program induced by QS-21 alone. (C) Heatmap displaying the relative expression levels of genes associated with macrophage activation and function across QS-21 and QS-21+QS-7 treatment conditions, illustrating broadly elevated expression in co-treated macrophages relative to QS-21 alone. (D) Gene Ontology (GO) enrichment analysis of biological processes significantly associated with differentially expressed genes in QS-21+QS-7-treated macrophages, with the top 12 significantly enriched processes ranked by FDR value. (E) Ingenuity Pathway Analysis (IPA) depicting significantly upregulated canonical pathways in QS-21-treated macrophages and the top 15 significantly upregulated canonical pathways in QS-21+QS-7-treated macrophages, ranked by activation Z-score. (F) Heatmap of IPA-derived activation Z-scores comparing the relative activation of canonical pathways associated with pro-inflammatory activation of macrophages and immunoregulatory signaling between QS-21 and QS-21+QS-7 treatment conditions. Data represent n = 3 independent biological replicates.

To further resolve the immunological pathways engaged by each treatment, Ingenuity Pathway Analysis (IPA) was performed on the differentially expressed genes from QS-21 and QS-21+QS-7 conditions. QS-21 treatment significantly altered 21 pathways (Benjamini-Hochberg corrected p-value; |Z-score| ≥ 2) (Supplementary Tables 9-10), of which 5 were significantly upregulated, including the NF1 Ras signaling pathway, breast cancer regulation by stathmin1, lung ionic balance signaling pathway, BBsome signaling pathway and response of EIF2AK1 (HRI) to heme deficiency (Figure 5E). In contrast, QS-21+QS-7 co-treatment significantly altered 71 pathways, of which a total of 41 pathways were upregulated (Supplementary Tables 11-12) and the top 15 pathways are shown (Figure 5E), reflecting a substantially expanded pathway activation landscape. Notably, the combination treatment engaged canonical pro-inflammatory and macrophage activation pathways—including HMGB1, IL-6, IL-17, TNF, and IL-1 signaling, as well as macrophage classical activation and Th1 pathways—at higher activation Z-scores than QS-21 alone (Figure 5F). Concurrently, pathway analysis predicted reduced activity of pathways associated with negative regulation of inflammatory activation, including PPAR, GADD45, RAR activation and FXR/RXR signaling, in QS-21+QS-7-treated macrophages, suggesting a coordinated shift toward a more sustained and less restrained pro-inflammatory state. Taken together, the transcriptomic data reinforce and extend the functional observations from surface marker and cytokine profiling, establishing that QS-7 potentiates the immunostimulatory activity of QS-21 at the transcriptional level by amplifying pro-inflammatory gene expression programs and pathway activation, while concurrently showing predicted downregulation of negative regulatory pathways.

## Discussion

Saponin-based adjuvants derived from *Q. saponaria* have emerged as potent immunostimulatory components of licensed vaccine formulations, with QS-21 representing the most extensively characterized fraction and a clinically validated constituent of the AS01 adjuvant system. Despite belonging to the same biological extract, the function and immunological properties, and toxicity of QS extracts other than QS-21 remain inadequately explored. The present study sought to address this gap through a systematic, side-by-side immunological characterization of QS-21 and QS-7 in the THP-1 human macrophage model, dissecting their relative potencies and functional profiles across multiple parameters of innate immune activation.

### Effects on cell viability

Cytotoxicity profiling established that QS-21 and QS-7 exhibit broadly comparable dose-dependent reductions in cellular metabolic activity in THP-1 macrophages across most of the tested concentration range, with QS-21 demonstrating greater cytotoxic potency at the highest tested concentration of 20 μg/mL. In contrast, QS-7 was considerably better tolerated than QS-21 in A549 epithelial cells. This finding shows that the difference in structural properties between QS-21 and QS-7, specifically the absence of a longer acyl chain in QS-7 and its higher hydrophilicity may be associated with a dampening of cytotoxicity. Further exploration of QS-extracts can help establish such structure-function correlations, which can assist in the design of new-to-nature adjuvants. An important limitation of the viability assessment is that PrestoBlue HS measures cellular metabolic activity as a proxy for viability rather than directly quantifying cell death. Although this assay provides a useful comparative assessment of the effects of QS-21 and QS-7 when administered individually, metabolic activity may not fully reflect treatment-associated cell death or cellular stress.

### Surface activation, cytokine secretion, and transcriptional responses

Flow cytometric analysis revealed that QS-21 robustly upregulated co-stimulatory and antigen-presentation molecules CD80, CD86, and HLA-DR in a dose-dependent manner, whereas QS-7 alone failed to drive significant surface activation marker expression across its full tested dose range. Notably, QS-7 significantly potentiated QS-21-mediated macrophage surface activation upon co-treatment in a dose-dependent manner—resulting in an immunogenic profile that was more accentuated than achievable by either adjuvant alone.

In contrast, multiplex cytokine and chemokine profiling demonstrated that QS-7 independently drives dose-dependent secretion of a broad pro-inflammatory mediator panel—including TNFα, IL-6, IL-1β, IL-18, GM-CSF, CXCL1, and CCL2—albeit at attenuated potency relative to QS-21 at equivalent concentrations. This dissociation between surface activation marker expression and cytokine secretion in the macrophage response to QS-7 suggests that these two arms of macrophage activation may be engaged through (at least partially) distinct upstream mechanisms, warranting further mechanistic investigation.

Transcriptional profiling by RNA sequencing further revealed that while QS-7 alone did not induce significant differential gene expression in our assays, its co-administration with QS-21 markedly broadened and amplified the pro-inflammatory transcriptional landscape, with pathway analysis predicting enhanced activity of canonical macrophage activation pathways—including HMGB1, IL-6, IL-17, TNF, and IL-1 signaling—relative to QS-21 alone, alongside predicted downregulation of immune regulatory pathways such as PPAR signaling, GADD45 signaling, RAR activation, and FXR/RXR signaling. These transcriptional findings reinforce and extend the functional observations from surface marker and cytokine profiling, establishing QS-7 as a biologically active immunomodulator whose co-administration with QS-21 consistently amplifies macrophage activation beyond levels attributable to QS-21 alone. However, further validation, including RT-qPCR analysis of selected differentially expressed genes would help confirm the RNA-seq findings and strengthen the interpretation of the pathways identified (Supplementary Table 13).

An important consideration when interpreting the immunological findings is the potential contribution of treatment-associated cytotoxicity. In the THP-1 model, both QS-21 and QS-7 reduced cellular metabolic activity at concentrations of 10 µg/mL and above, which overlap with concentrations used in the flow cytometry and cytokine profiling experiments. Consequently, it cannot be excluded that cellular stress or damage at these concentrations contributed, at least in part, to the observed inflammatory responses.

### Potential structural basis for differential immunomodulatory activity

The structural distinctions between QS-21 and QS-7 offer a plausible basis for the differential and complementary activation profiles observed in this study. Previous studies have linked the long, chiral acyl side chain of QS-21 to interactions with cholesterol-rich membrane domains, cholesterol-dependent endocytosis, lysosomal accumulation, and membrane destabilization. These events have been associated with Syk kinase and cathepsin B-dependent signaling upstream of NF-κB and inflammasome activation in human APCs. In contrast, the truncation of this acyl domain to a short acetyl group in QS-7, together with the addition of two extra monosaccharide residues that increase its overall hydrophilicity, may reduce its capacity for membrane intercalation (11). Because QS-21 and QS-7 differ in both acyl substitution and oligosaccharide composition, this comparison can’t isolate the contribution of either feature to the biological activity. Studies using structurally defined analogues that vary these features individually would help distinguish their effects on membrane interactions, cell viability, and immunomodulatory responses. Whether these structural differences account for the limited surface marker and transcriptional responses observed with QS-7 remains to be established. The apparent dissociation between QS-7’s limited effect on surface marker expression and its capacity to independently induce cytokine and chemokine secretion further suggests that these responses may involve mechanisms distinct from those associated with QS-21. One possibility is that the extended oligosaccharide domain of QS-7 contributes to signaling through C-type lectin receptors on macrophages, potentially providing an input that is less dependent on membrane perturbation and lysosomal signaling. However, this remains speculative, as receptor engagement and downstream signaling were not directly assessed.

Within this framework, we propose as a working hypothesis that the potentiating effect of QS-7 on QS-21—driven macrophage activation may arise from the convergence of structurally distinct inputs: membrane and lysosomal perturbation associated with QS-21, together with a potential additional signaling contribution from QS-7. Their combined effects may also be associated with altered macrophage lipid homeostasis. Consistent with this possibility, pathway analysis predicted reduced activity of PPAR, RAR, and FXR/RXR signaling under combination treatment—pathways that are involved in lipid and sterol sensing and the regulation of inflammatory responses. Direct assessment of this hypothesis, for example through cholesterol depletion, cathepsin B inhibition, or lectin receptor blockade in the context of QS-7 and QS-21 co-treatment, represents an important avenue for future mechanistic work. Of particular translational significance was the emergence of IL-12p70 and IFNγ secretion exclusively under co-treatment conditions. However, given that many replicate values were below the LLOQ, these findings should be interpreted with caution. While IL-12p70 is associated with Th1-polarizing immune responses relevant to QS saponin-based vaccine formulations, the biological significance of IFN-γ secretion in this THP-1-derived macrophage model requires further validation.

### Implications and future directions

These findings carry meaningful implications for the rational design of next-generation adjuvant systems. Specifically, the observed capacity of QS-7 to enhance the immunostimulatory activity of QS-21 suggests that these two saponin fractions may have complementary effects when combined. Whether this enhanced immunological response could permit a reduction in the effective dose of QS-21 remains to be determined. Future studies using a concentration matrix of QS-21, both alone and in combination with QS-7, will be required to establish whether such a dose-sparing effect can be achieved and whether it translates into reduced hemolytic toxicity. These findings identify QS-7 as a potentially complementary adjuvant partner in QS-21-based formulations, warranting further investigation of its effects on immunological activity, dose requirements, and tolerability.

The present study opens several avenues for future investigation. The absence of significant differential gene expression with QS-7 alone at the specific experimental conditions and analytical thresholds used in this study, precluded pathway-level mechanistic resolution of its independent immunomodulatory activity; future transcriptional studies employing other QS-7 concentrations and treatment timepoints may yield sufficient signal to delineate the gene networks independently engaged by QS-7. Validation of the findings reported here in primary human monocyte-derived macrophages and dendritic cells would further strengthen their translational relevance beyond the THP-1 model. The cooperative immunostimulatory activity observed for the QS-21+QS-7 combination also warrants direct comparison with the clinically established AS01 adjuvant system, in which QS-21 is combined with the TLR4 agonist MPLA. Direct biophysical characterization of QS-7’s membrane and cholesterol interactions, relative to QS-21, would help substantiate the structural basis proposed here for their differential and cooperative mechanisms of macrophage activation. Finally, *in vivo* evaluation of antigen-specific immune responses elicited by QS-21+QS-7 co-formulations, and assessment of their compatibility with liposomal-based delivery platforms, will be essential to fully realize the translational potential suggested by these findings.

## Conclusion

QS-21 and QS-7 exhibited distinct immunomodulatory profiles in THP-1 derived macrophages. QS-21 induced stronger surface activation and transcriptional responses, whereas QS-7 promoted cytokine and chemokine secretion despite limited effects on surface markers and transcription at the timepoints examined. Co-treatment enhanced several measures of macrophage activation, supporting further investigation of their combined activity. This study establishes a multimodal workflow for characterizing the viability and immunomodulatory effects of saponins providing a basis for investigating structure-function relationships. Together with AI-driven design and synthetic biology pipelines, such approaches could support the development of a diverse library of immunomodulators with tailored properties.

## Acknowledgements

We thank the Technology Center for Genomics & Bioinformatics (TCGB) at UCLA for providing sequencing and bioinformatics services and data generation. We thank Peter H. Winegar for providing us with the saponin chemical structures. GPT-5.6 Luna was used to assist with the development, debugging and refinement of R code used in this study. We thank Xixi Zhao for careful review and assessment of QS-21 purity. All AI-generated code was reviewed, modified as necessary, and validated by the authors. This publication was supported by the National Institute of Allergy and Infectious Diseases (NIAID) of the National Institutes of Health (5R01AI186111-03). The content is solely the responsibility of the authors and does not necessarily represent the official views of the National Institutes of Health.

## Bibliography

1. Molina GI. Saggio sulla Storia Naturalle deChile. 1782.

2. Kensil CR, Patel U, Lennick M, Marciani D. Separation and characterization of saponins with adjuvant activity from Quillaja saponaria Molina cortex. J Immunol. 1991 Jan 15;146(2):431–7.

3. Soltysik S, Wu J-Y, Recchia J, Wheeler DA, Newman MJ, Coughlin RT, et al. Structure/function studies of QS-21 adjuvant: assessment of triterpene aldehyde and glucuronic acid roles in adjuvant function. Vaccine. 1995 Jan;13(15):1403–10.

4. Press JB, Reynolds RC, May RD, Marciani DJ. Structure/function relationships of immunostimulating saponins. Bioactive natural products (part E). Elsevier; 2000. p. 131–74.

5. Talukdar P, Winegar PH, Hudson GA, Astolfi MCT, Inman JL, Keasling JD, et al. From bark to bench: innovations in QS-21 adjuvant characterization and manufacturing. Front Immunol. 2025 Dec 2;16:1677995.

6. Welsby I, Detienne S, N’Kuli F, Thomas S, Wouters S, Bechtold V, et al. Lysosome-Dependent Activation of Human Dendritic Cells by the Vaccine Adjuvant QS-21. Front Immunol. 2016;7:663.

7. Chea EK, Fernández-Tejada A, Damani P, Adams MM, Gardner JR, Livingston PO, et al. Synthesis and preclinical evaluation of QS-21 variants leading to simplified vaccine adjuvants and mechanistic probes. J Am Chem Soc. 2012 Aug 15;134(32):13448–57.

8. Liu G, Anderson C, Scaltreto H, Barbon J, Kensil CR. QS-21 structure/function studies: effect of acylation on adjuvant activity. Vaccine. 2002 Jun 21;20(21–22):2808–15.

9. Deng K, Adams MM, Gin DY. Synthesis and structure verification of the vaccine adjuvant QS-7-Api. Synthetic access to homogeneous Quillaja saponaria immunostimulants. J Am Chem Soc. 2008 May 7;130(18):5860–1.

10. Kensil CR. Saponins as vaccine adjuvants. Crit Rev Ther Drug Carrier Syst. 1996;13(1–2):1–55.

11. Kensil CR, Wu JY, Anderson CA, Wheeler DA, Amsden J. QS-21 and QS-7: purified saponin adjuvants. Dev Biol Stand. 1998;92:41–7.

12. Kensil CA. Saponin adjuvant compositions. U.S; 6231859, 2021.

13. Stylianou VV, Bertram KM, Van Anh Vo, Dunn EB, Baharlou H, Terre DJ, et al. Innate immune cell activation by adjuvant AS01 in human lymph node explants is age independent. The Journal of Clinical Investigation. 2024 Nov 15;

14. Detienne S, Welsby I, Collignon C, Wouters S, Coccia M, Delhaye S, et al. Central Role of CD169+ Lymph Node Resident Macrophages in the Adjuvanticity of the QS-21 Component of AS01. Sci Rep. 2016 Dec 20;6:39475.

15. Liu Y, Zhao X, Gan F, Chen X, Deng K, Crowe SA, et al. Complete biosynthesis of QS-21 in engineered yeast. Nature. 2024 May 8;629(8013):937–44.

16. Ragupathi G, Damani P, Deng K, Adams MM, Hang J, George C, et al. Preclinical evaluation of the synthetic adjuvant SQS-21 and its constituent isomeric saponins. Vaccine. 2010 Jun 11;28(26):4260–7.

17. Talukdar P. RNA-seq Data Analysis Scripts for Differential Gene Expression and Functional Enrichment Analysis [Computer software]. Zenodo. [Internet]. Zenodo; 2026 [cited 2026 Oct 31]. Available from: 10.5281/zenodo.23850488

18. Blighe K, Rana S, Lewis M. EnhancedVolcano: Publication-ready volcano plots with enhanced colouring and labeling. [Internet]. 2018 [cited 2026 Jul 16]. Available from: https://github.com/kevinblighe/enhancedvolcano

19. Larsson J. eulerr: Area-Proportional Euler Diagrams with Ellipses. 2018;

20. Wickham H. Elegant Graphics for Data Analysis. New York: Springer-Verlag; 2016.

21. Smith CL, Richardson B, Rubsamen M, Cameron MJ, Cameron CM, Canaday DH. Adjuvant AS01 activates human monocytes for costimulation and systemic inflammation. Vaccine. 2024 Jan 12;42(2):229–38.

22. Marty-Roix R, Vladimer GI, Pouliot K, Weng D, Buglione-Corbett R, West K, et al. Identification of QS-21 as an Inflammasome-activating Molecular Component of Saponin Adjuvants. J Biol Chem. 2016 Jan 15;291(3):1123–36.

23. Afkarian M, Sedy JR, Yang J, Jacobson NG, Cereb N, Yang SY, et al. T-bet is a STAT1-induced regulator of IL-12R expression in naïve CD4+ T cells. Nat Immunol. 2002 Jun;3(6):549–57.

24. Henry CJ, Ornelles DA, Mitchell LM, Brzoza-Lewis KL, Hiltbold EM. IL-12 produced by dendritic cells augments CD8+ T cell activation through the production of the chemokines CCL1 and CCL17. J Immunol. 2008 Dec 15;181(12):8576–84.

25. Wenner CA, Güler ML, Macatonia SE, O’Garra A, Murphy KM. Roles of IFN-γ and IFN-α in IL-12-induced T helper cell-1 development. The Journal of Immunology. 1996 Feb 15;156(4):1442–7.

26. Ip WKE, Hoshi N, Shouval DS, Snapper S, Medzhitov R. Anti-inflammatory effect of IL-10 mediated by metabolic reprogramming of macrophages. Science. 2017 May 5;356(6337):513–9.

27. Awasthi A, Carrier Y, Peron JPS, Bettelli E, Kamanaka M, Flavell RA, et al. A dominant function for interleukin 27 in generating interleukin 10-producing anti-inflammatory T cells. Nat Immunol. 2007 Dec;8(12):1380–9.

