## Supplemental Figures for "Differential and Cooperative Immunomodulation by Saponin Adjuvants QS-21 and QS-7 in Human THP-1 Macrophages"

**Supplementary Figure 1: Flow cytometric analysis of surface marker expression in THP-1 cells.**

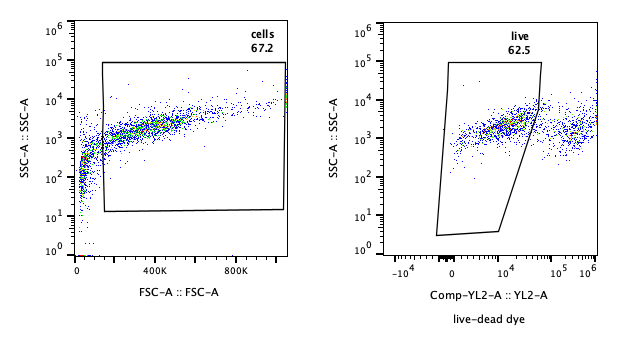

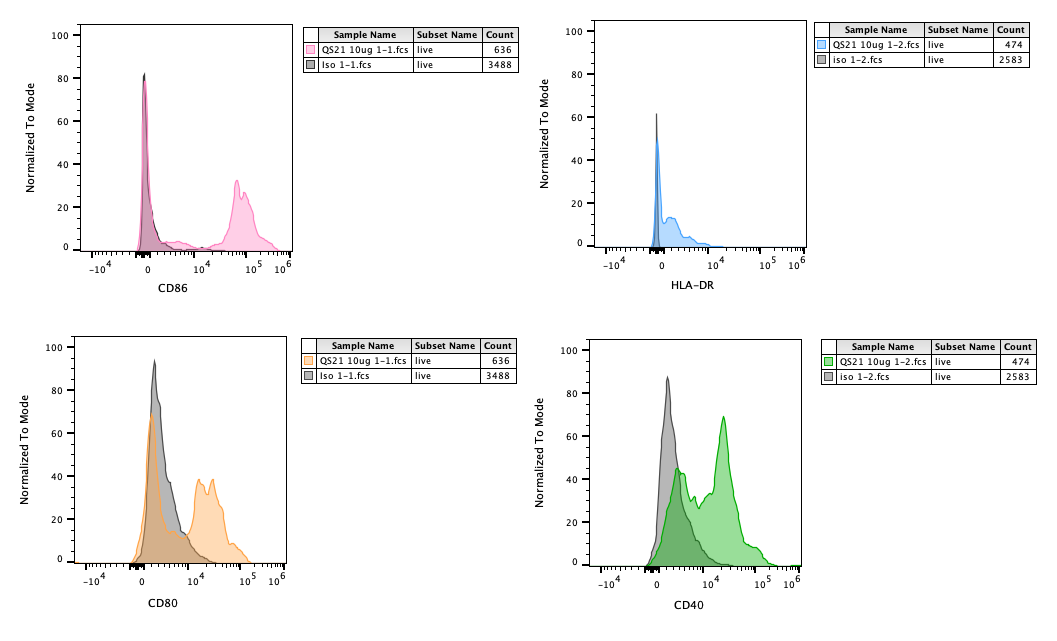

**Supplementary Figure 1:** Representative flow cytometry gating plots showing THP-1 derived macrophages were first gated based on forward scatter (FSC) and side scatter (SSC), followed by gating of viable cells using the Live/Dead Fixable Red viability dye. Representative flow cytometry histograms showing the expression of CD86, HLA-DR, CD80 and CD40 in THP-1 cells following QS21 treatment (10 µg/mL). Coloured histograms represent cells stained with the respective fluorescently labelled antibodies, while grey histograms represent the corresponding isotype controls. All analyses were performed on the gated live-cell population. The geometric mean fluorescence intensity (gMFI) of each marker was determined using the corresponding isotype control as a reference for background fluorescence.

**Supplementary Figure 2: Absolute cytokine/chemokine concentrations post-treatment with QS-21, QS-7 or QS-21+QS-7.**

(A) **Absolute Cytokine/Chemokine Concentrations for QS-21 Treatment**

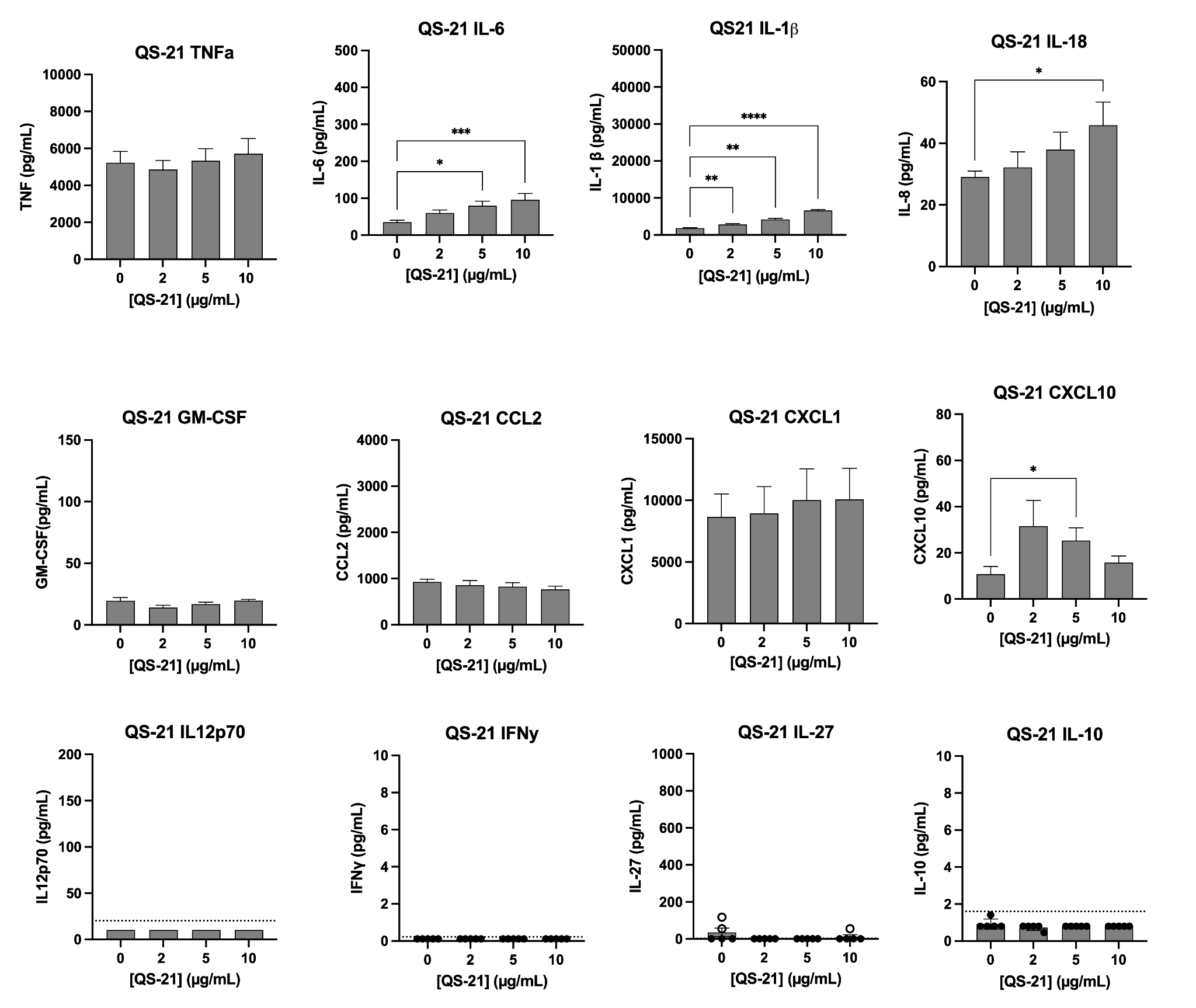

**(B) Absolute Cytokine/Chemokine Concentrations for QS-7 Treatment**

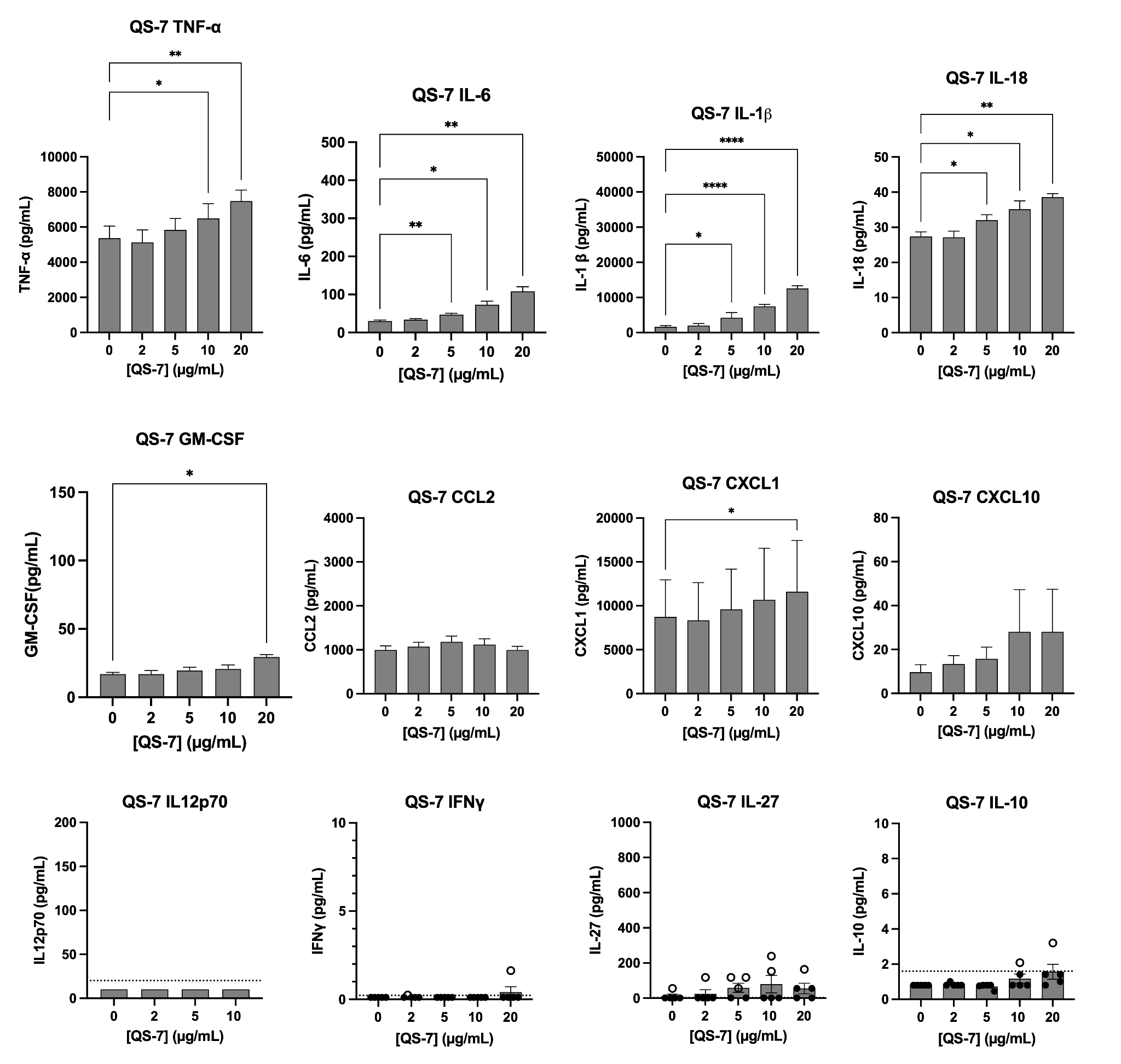

**(C) Absolute Cytokine/Chemokine Concentrations for QS-21+QS-7 Treatment**

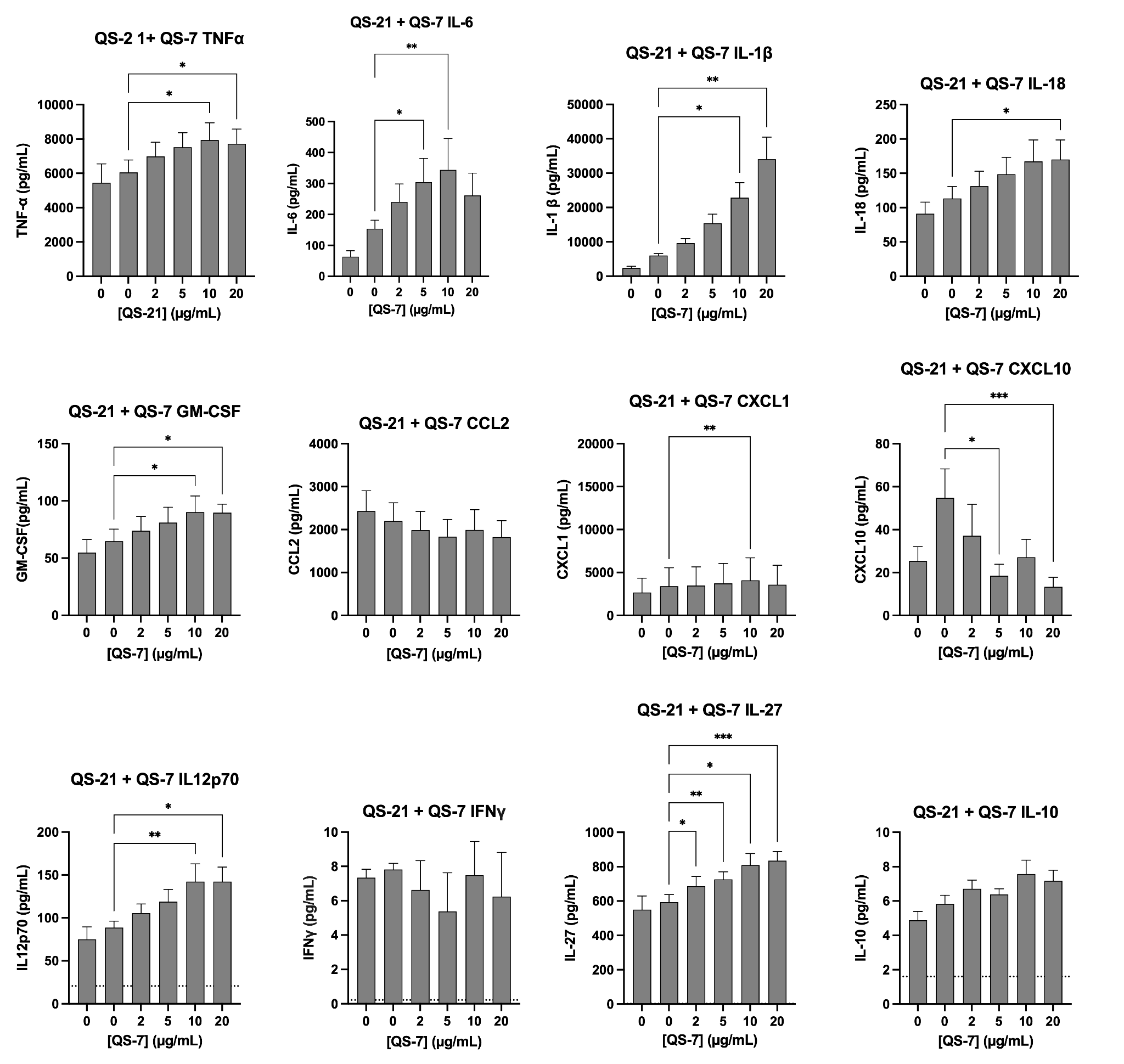

**Supplementary Figure 2:** Absolute cytokine/chemokine concentrations post-treatment with QS-21, QS-7 or QS-21+QS-7. Luminex multiplex quantification of cytokine and chemokine secretion in THP-1 macrophages treated with increasing concentrations of QS-21 or QS-7 alone, or QS-7 in combination with a fixed sub-maximal concentration of QS-21 (5 μg/mL). Data are presented as mean ± SEM of n = 5 independent biological replicates for QS-21 and QS-7 alone treatments. Data are presented as mean ± SEM of n = 5 independent biological replicates for QS-21+QS-7 combination treatments. For IL-12p70, IFNγ, IL-10 and IL-27 many replicate values when the cells were treated with QS-21 or QS-7 alone, were below the limit of detection (LOD). The LOD is represented by a dashed line. Replicate values above the LOD are represented with an open circle and those below are represented as a filled circle for the purpose of visualization only. For QS-21+QS-7 combination treatment, values for these four analytes were above the LOD.

**Supplementary Figure 3: PCA plot from RNA sequencing data.**

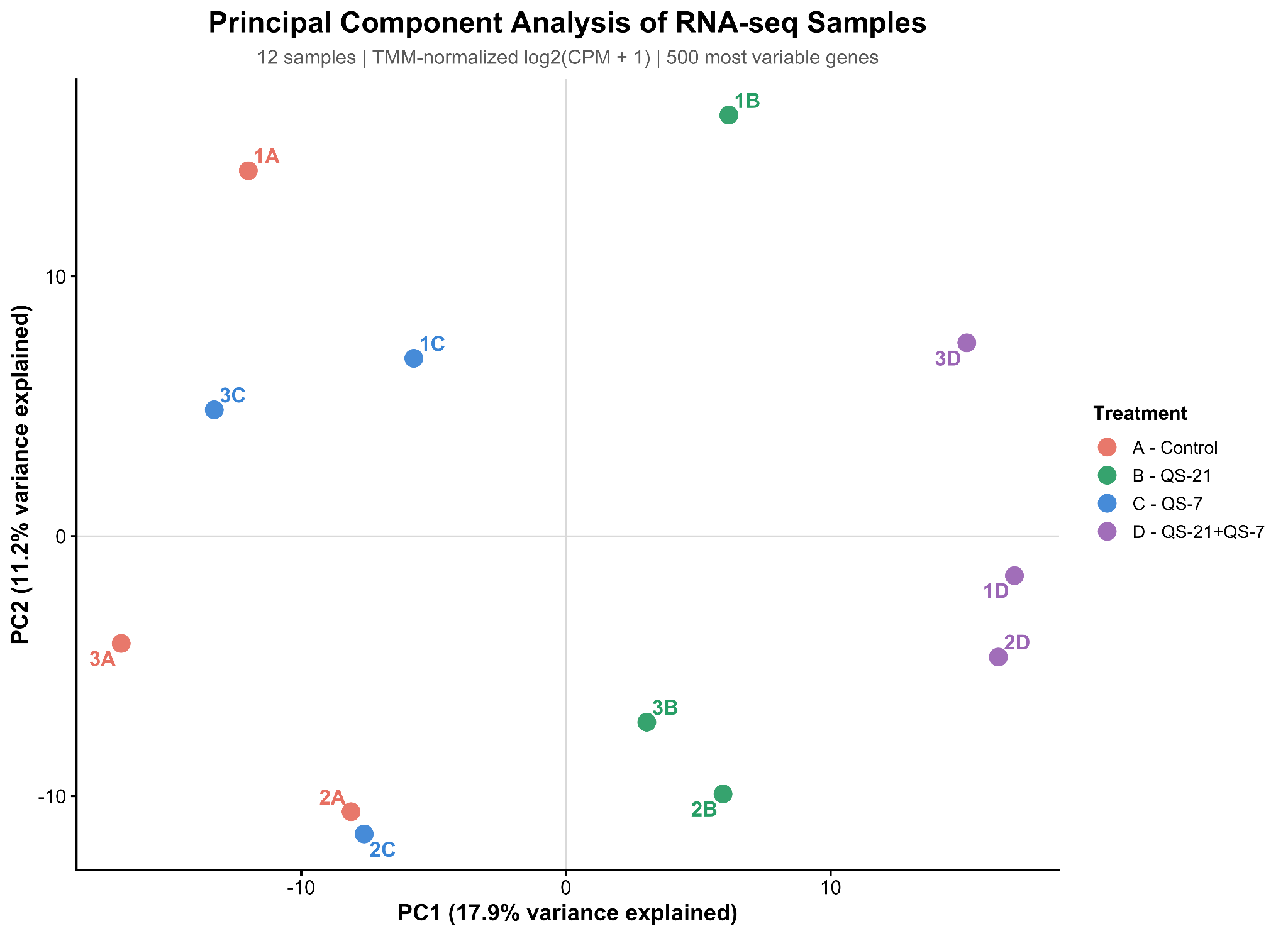

**Supplementary Figure 3:** PCA plots from RNA sequencing data of THP-1 macrophages treated with QS-21, QS-7 or QS21+QS-7.: A, B, C and D represent the different treatments (and control) and 1,2 and 3 represent biological replicates.
