## Supplemental Table 13 for "Differential and Cooperative Immunomodulation by Saponin Adjuvants QS-21 and QS-7 in Human THP-1 Macrophages"

**Supplemental Table 13. Cross-platform comparison of transcriptional, cytokine secretion, and cell-surface activation responses to QS-21 and QS-7 in THP-1-derived macrophages.**

| **Treatment (dose)** | **RNA-seq, 5 h** | **Cytokines, 24 h** | **Surface markers, 24 h** | **Interpretation** |
| --- | --- | --- | --- | --- |
| QS-21 (5 ug/mL) | 16 upregulated, 2 downregulated DEGs | Significant TNFα, IL-6, IL-1β, IL-18, CXCL1, CXCL10, CCL2 | Significant CD86; CD80, CD40, HLA-DR not significant | Broad inflammatory activity across platforms, with selective surface-marker activation |
| QS-7 (5 ug/mL) | No significant DEGs | Significant CXCL1 and CCL2; other measured cytokines not significant | No significant changes in CD80, CD86, CD40, HLA-DR | Selective chemokine secretion despite no significant early DEGs or surface-marker changes |
| QS-21 + QS-7 (5 ug/mL each) | 61 upregulated, 8 downregulated DEGs | No significant enhancement over QS-21 alone reported at 5 + 5 µg/mL | No significant enhancement over QS-21 alone reported at 5 + 5 µg/mL | Broader early transcriptional signature relative to control without corresponding significant enhancement of 24-hour protein responses |

**Supplemental Table 13. Cross-platform comparison of transcriptional, cytokine secretion, and cell-surface activation responses to QS-21 and QS-7 in THP-1-derived macrophages.** Summary of immunological responses following treatment with QS-21 (5 µg/mL), QS-7 (5 µg/mL), or their combination (5 µg/mL each). RNA sequencing was performed at 5 hours to characterize early transcriptional responses, whereas cytokine and chemokine secretion (Luminex) and cell-surface activation marker expression (flow cytometry) were assessed at 24 hours. Differentially expressed genes (DEGs) were defined using an FDR < 0.05 and |log2FC| ≥ 2. The table highlights areas of agreement and divergence across platforms. Differences in sampling time, assay sensitivity, transcriptional and protein regulation, and protein accumulation may contribute to the observed discrepancies. Notably, combination treatment produced a broader early transcriptional response without significant enhancement of cytokine secretion or surface-marker expression relative to QS-21 alone at matched concentrations. Enhanced functional responses were primarily observed at higher QS-7 concentrations, indicating concentration-dependent effects.
