## Supplemental Table 2 for "Differential and Cooperative Immunomodulation by Saponin Adjuvants QS-21 and QS-7 in Human THP-1 Macrophages"

**Supplementary Table 2: Flow Cytometry gMFI of replicates**

QS-21 Flow cytometry gMFI of each replicate: CD80

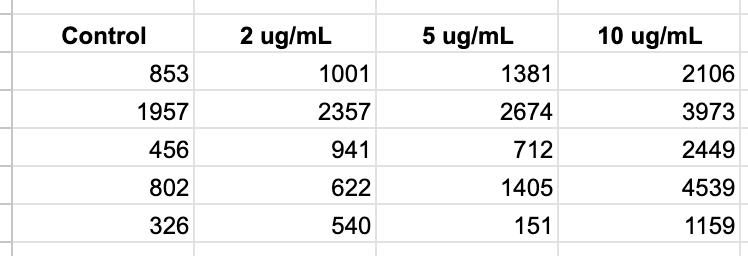

QS-21 Flow cytometry gMFI of each replicate: CD86
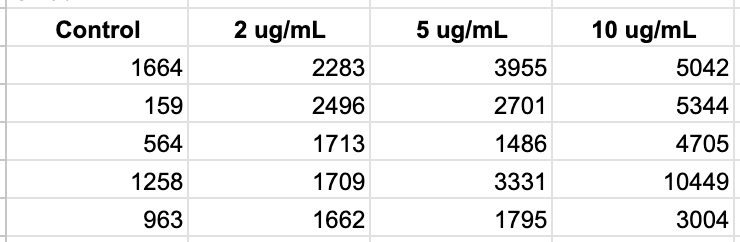

QS-21 Flow cytometry gMFI of each replicate: CD40
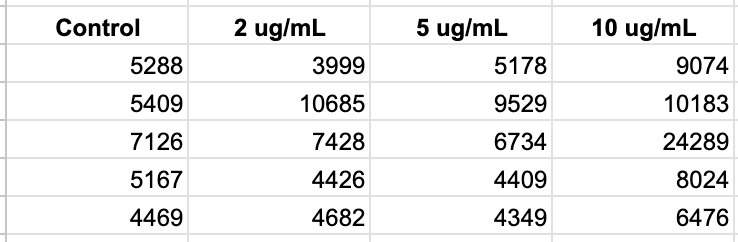

QS-21 Flow cytometry gMFI of each replicate: HLA-DR

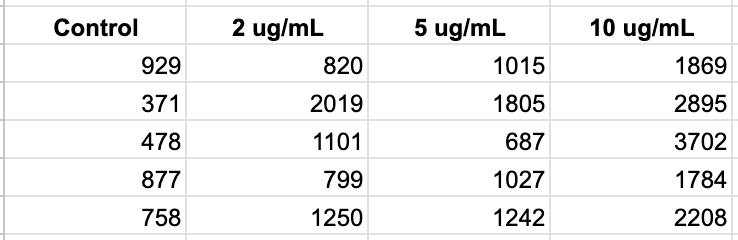

QS-7 Flow cytometry gMFI of each replicate: CD80

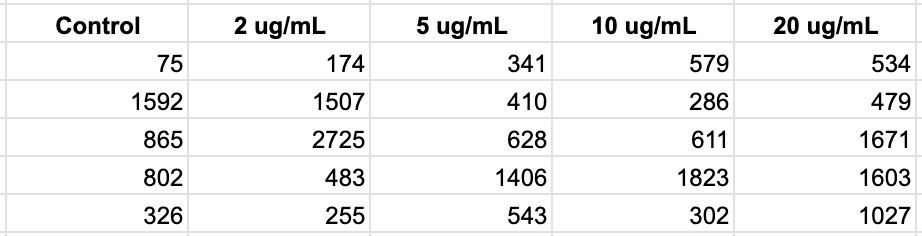

QS-7 Flow cytometry gMFI of each replicate: CD86
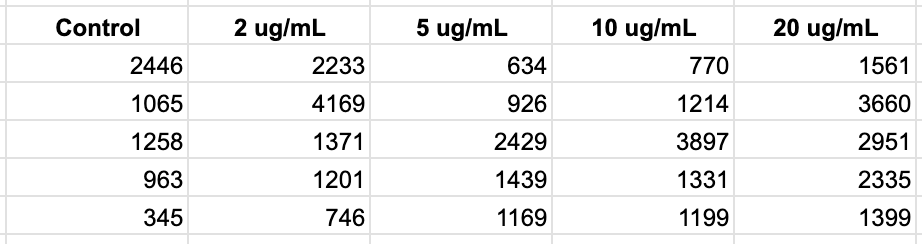

QS-7 Flow cytometry gMFI of each replicate: CD40

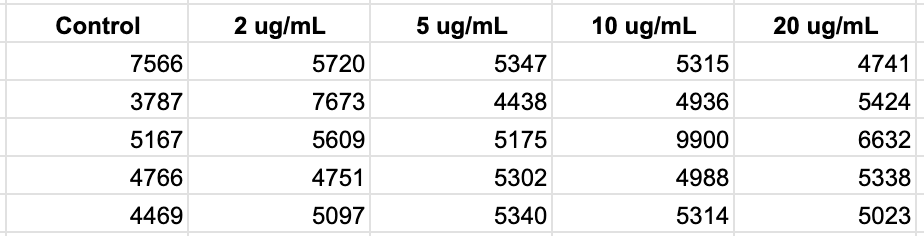

QS-7 Flow cytometry gMFI of each replicate: HLA-DR

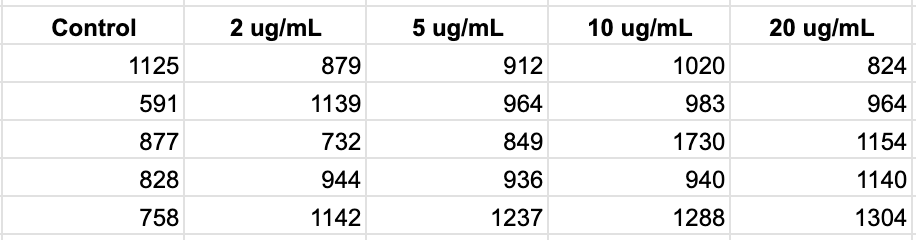

QS-21+QS-7 flow cytometry MFI of each replicate: CD80

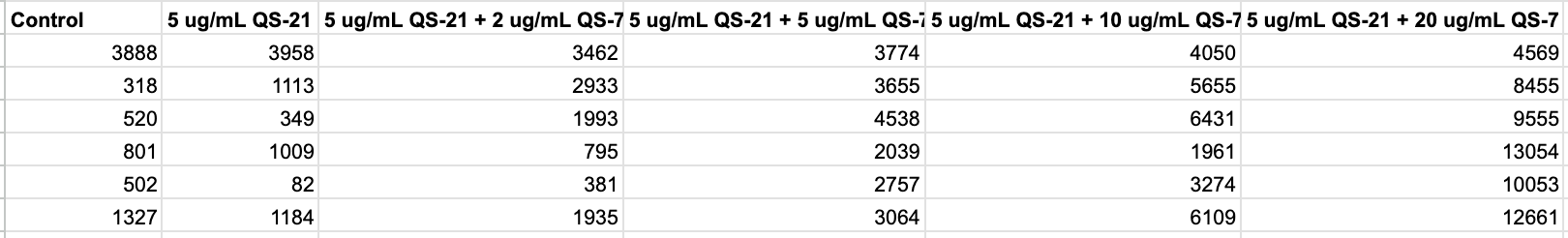

QS-21+QS-7 flow cytometry MFI of each replicate: CD86

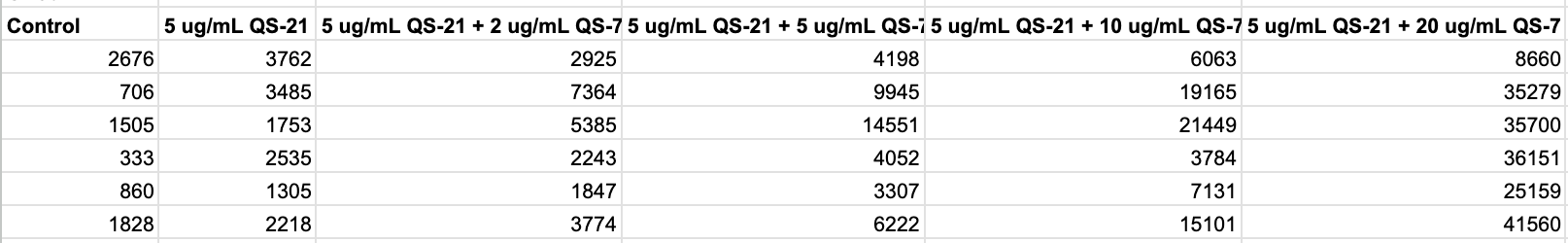

QS-21+QS-7 flow cytometry MFI of each replicate: CD40

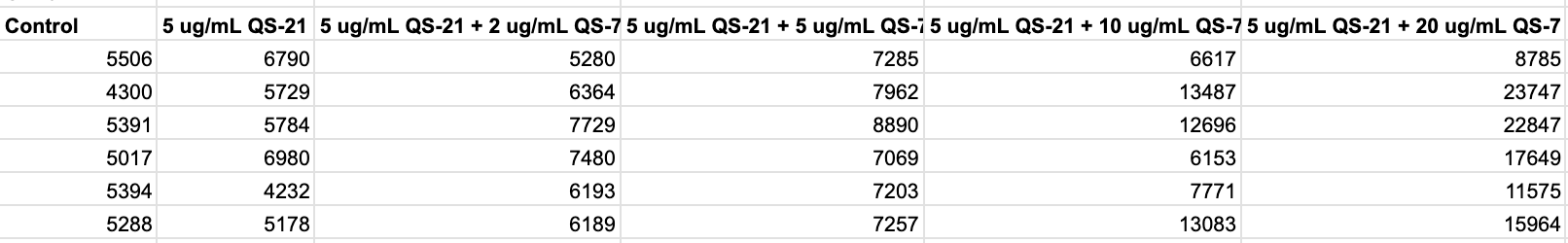

QS-21+QS-7 flow cytometry MFI of each replicate: HLA-DR

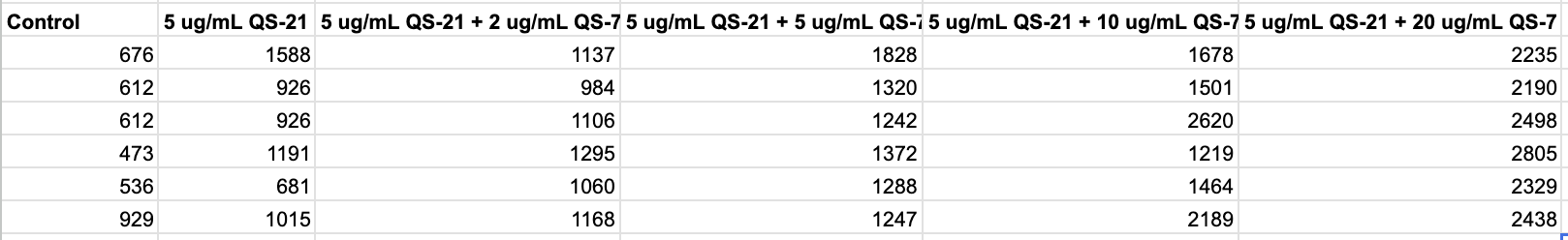
