## Supplemental Table 9 for "Differential and Cooperative Immunomodulation by Saponin Adjuvants QS-21 and QS-7 in Human THP-1 Macrophages"

**Supplementary Table 9: IPA generated canonical pathways with significant differential expression in QS-21 treated THP-1 macrophages**

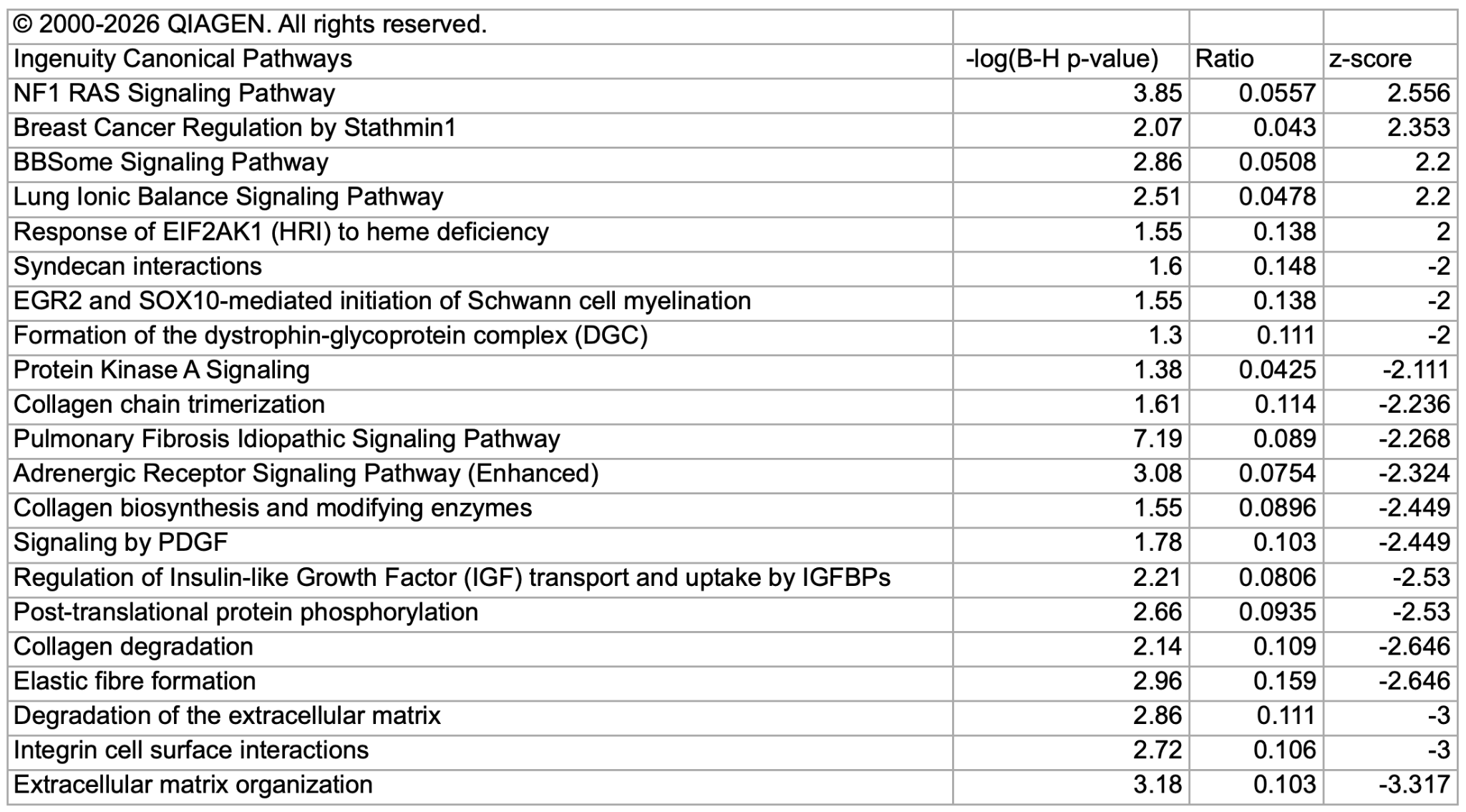
